# Spontaneous binding of single-stranded RNAs to RRM proteins visualised by unbiased atomistic simulations with rescaled RNA force field

**DOI:** 10.1101/2022.07.22.501120

**Authors:** Miroslav Krepl, Pavlina Pokorna, Vojtech Mlynsky, Petr Stadlbauer, Jiri Sponer

## Abstract

Recognition of single-stranded RNA (ssRNA) by RNA recognition motif (RRM) domains is an important class of protein-RNA interactions. Many such complexes were characterized using NMR and/or X-ray crystallography techniques, revealing ensemble-averaged pictures of the bound states. However, it is becoming widely accepted that better understanding of protein-RNA interactions would be obtained from ensemble descriptions. Indeed, earlier molecular dynamics (MD) simulations of bound states indicated visible dynamics at the RNA-RRM interfaces. Here, we report the first atomistic simulation study of spontaneous binding of short RNA sequences to RRM domains of HuR and SRSF1 proteins. Using millisecond-scale aggregate ensemble of unbiased simulations we were able to observe a few dozens of binding events. The HuR RRM3 utilizes a pre-binding state to navigate the RNA sequence to its partially disordered bound state and then to dynamically scan its different binding registers. The SRFS1 RRM2 binding is more straightforward but still multiple-pathway. The present study necessitated development of a goal-specific force-field modification scaling down the intramolecular vdW interactions of the RNA which also improves description of the RNA-RRM bound state. Our study opens a new avenue for large-scale atomistic investigations of binding landscapes of protein-RNA complexes and future perspectives of such research are discussed.

## Introduction

From synthesis to degradation, RNA molecules are almost constantly bound to proteins *in vivo*, forming ribonucleoprotein complexes. (1-5) The interactions are very variable and dynamical and a single RNA transcript binds to numerous proteins within its lifetime. (6) Understanding principles and function of protein-RNA recognition is therefore a cornerstone of RNA biology and a key component of many biological processes. Individual proteins can specialize on binding RNA single strands (ssRNA) as well as structured RNAs such as duplexes or other recurrent motifs. (2,7-9) Bases within ssRNA are fully exposed for a readout which is commonly utilized by proteins whose biological function involves targeted response towards specific RNA sequences. (10) Among such proteins, the RNA recognition motif (RRM) domain is the most prominent, being the most wide-spread RNA binding motif in eukaryotes. (11) Individual RRM domains can recognize various RNA sequences despite sharing a very conserved fold in which two α-helices are packed against an antiparallel β-sheet surface. (12) The partially exposed β-sheet surface is a canonical RNA-binding RRM site. However, other parts of the RRM, like the α-helices, can also be utilized for RNA binding. (12) Specificity of RRMs towards different RNA sequences can be astonishingly tuned by subtle variations of their surface-exposed amino acids. (13-17) The basic principles of RNA-RRM interactions were elucidated by NMR and/or X-ray crystallography techniques. However, complete understanding of the RNA-RRM recognition would require information extending beyond the static ensemble-averaged picture of the bound state, i.e., description of the RNA-RRM interactions as dynamic ensemble of conformations, which form with different populations and lifetimes. Knowledge of the complete free-energy binding landscape would allow unraveling how the proteins search for their RNA targets in the cellular pool of biomolecules and how different RNA sequences compete with each other for the binding.

Molecular dynamics (MD) simulations are a method for describing the Boltzmann distribution (populations) of molecules using carefully calibrated empirical potentials, commonly known as the force fields (*ff*s). MD has been successfully applied many times to study nucleic acid systems, including the protein-RNA complexes. (18-20) MD provides potentially infinite spatial and temporal resolution unrivalled by any experimental method and can significantly expand upon the information obtained from ensemble- and time-averaged experimental structural data. However, in practice, MD studies are also affected by fundamental limitations – the quality of the *ff* and the affordable timescale (sampling). The two issues are interconnected as *ff* inaccuracies are routinely discovered once longer timescales become computationally affordable. (18) Notorious problem of biomolecular *ff*s designed to simulate folded states is excessive compaction of unstructured ensembles. (21) This makes the description of ssRNAs challenging, as illustrated by studies of RNA tetranucleotides (TNs) (18,22-26) and tetraloops. (27-31) In case of TNs, the standard AMBER RNA *ff* leads to large populations of compact intercalated structures which is not in agreement with experiments. (32-34) For TLs, the excessive interactions in the unfolded states frustrate the free-energy landscape and hinder folding. The solution to this problem is not trivial as straightforwardly tuning the *ff* for a better description of small systems often deteriorates description of larger folded RNAs. (26) Possible solution is introducing additional *ff* terms orthogonal to the currently used parameters, such as the gHBfix potential which tunes the strength of hydrogen bonds. (26,35,36) Non-bonded-fix (NBfix) correction, a pair-specific adjustment of Lennard-Jones (LJ) combination rules, is another common approach to increase the parameterization flexibility. (30,37,38) An alternative option is to abandon the daunting task of a general *ff* reparametrization and instead adjust the parameters in a goal-directed manner for specific systems/processes. (39-42) This strategy is common in coarse-grained (CG) modelling. (18)

The prime goal of our current work was to capture spontaneous binding events of ssRNAs to their binding sites on the surface of RRMs, to complement the simple two-state picture of binding by visualization of representative number of continuous binding pathways. However, we found out that it is virtually impossible to observe any binding events with the standard atomistic RNA OL3 AMBER *ff (43)* that is designed to simulate folded RNAs. The free ssRNAs rapidly and irreversibly collapsed due to RNA self-interactions, preventing the binding. Thus, we had to develop a suitable *ff* modification that reduces the spurious RNA self-interactions. We directly address this problem via NBfix by rescaling the well-depth value (ε) of the Lennard-Jones (LJ) potentials between selected RNA-RNA atomic pairs. We abbreviate the new modification as stafix (<u>sta</u>cking <u>fix</u>) as it is aimed at weakening the stacking and vdW interactions as opposed to hydrogen bonds.

Stafix entirely eliminated occurrence of the spurious ssRNA self-interactions and allowed us to perform almost a millisecond of standard (unbiased) MD simulations where we documented few dozens of spontaneous binding events of short ssRNA sequences to two different RRM proteins – the HuR RRM3 and SRSF1 RRM2. The RRM3 domain of HuR (44,45) (Figure 1a) recognizes either U-rich or AU-rich segments of mRNA in a canonical mode typical for RRM domains (11) and its crystal structure showed it can bind four or five nucleotides. (44) However, a large degree of disorder was detected at its protein-RNA interface, with only pockets p2 and p3 being always resolved within the asymmetric unit of the X-ray structure. (44) In contrast, the SRSF1 RRM2 recognizes GGA triplet motif in mRNA via well-defined binding pockets. (46) Here, we for the first time sample the complete binding funnel of both complexes and conclude that differences between the native protein-RNA interfaces are already reflected in details of their binding pathways. Namely, the HuR RRM3 utilizes a previously unknown ‘pre-binding mode’ which navigates the binding and then facilitates fast binding register exchanges without fully separating the RNA from the protein. By separating the register exchange (i.e. sliding) from the native binding mode, the HuR RRM3 might be able to efficiently scan the RNA for its target sequence. In contrast, the SRSF1 RRM2 does not seem to utilize any pre-binding mode and displays very slow binding register exchanges.

**Figure 1.**
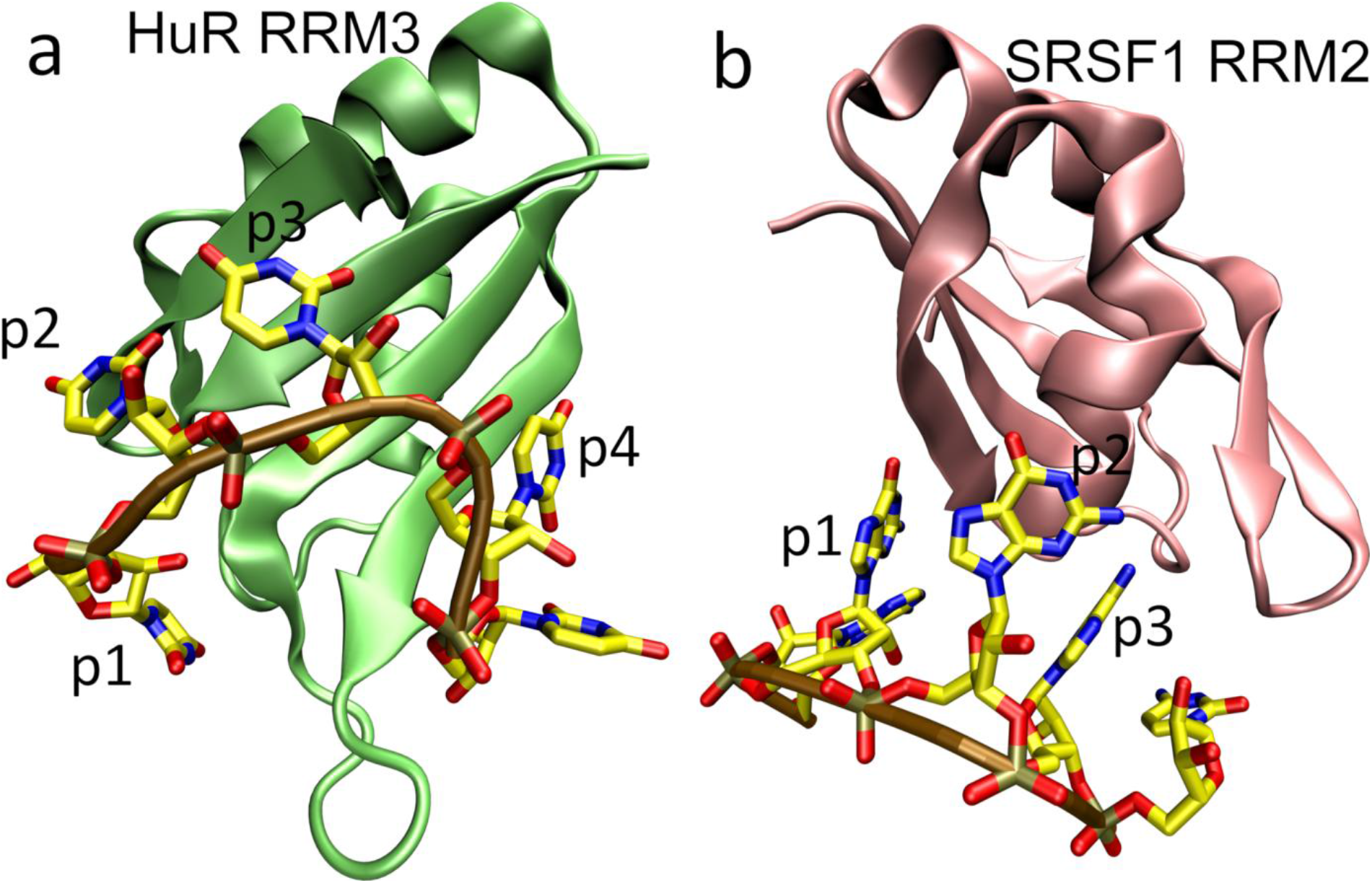
Protein-RNA complexes studied in this work – HuR RRM3 and SRSF1 RRM2. (44,46) a) 3D representation of the HuR RRM3 (lime) complexed with 5′-UUUUU-3′ RNA. Nitrogen, carbon, oxygen and phosphorus atoms in the RNA are in blue, yellow, red, and brown, respectively. The RNA backbone is indicated by ochre tube. Pockets p2 and p3 recognize only uracils and are well-defined whereas pockets p1 and p4 show significant degree of disorder and can also accept adenines. b) SRSF1 RRM2 (pink) complexed with 5′-AGGAC-3′ RNA. All three pockets are well-defined.

The stafix modification was absolutely essential to execute our study as otherwise the RNA molecules strongly prefer binding-incompetent spurious self-interactions rather than sampling potential binding sites on the proteins’ surfaces. Even if real ssRNAs sample to certain extent the compact structures (47) which we prevent by stafix, they would be likely off-pathway intermediate states with respect to the binding process. Thus, stafix is justifiable and should not fundamentally distort the studied process. Importantly, the stafix also significantly improves simulations of fully-bound protein-ssRNA complexes, which further substantiates our approach.

In summary, we provide the first atomistic insight into the complex multiple-pathway processes of spontaneous binding of ssRNA to the RRM proteins and suggest that RRM proteins utilize very diverse binding mechanisms to communicate with their RNA targets.

## Materials and methods

### Selection of initial structures

We used X-ray structure of the HuR RRM3 protein-RNA complex (PDB: 6GC5) (44) as the starting structure for simulations with either 5′-UUUUU-3′ (chains A and E) or 5′-UUUA-3′ (chains C and G) RNAs bound. Coordinates of the U_1_ nucleotide in the 5′-UUUUU-3′ system were obtained from chain G after first aligning the protein chains A and C to overlap. Coordinates of the non-target 5′-CCCCC-3′ RNA were obtained by mutating the 5′-UUUUU-3′ sequence. Structure of the free HuR RRM3 was prepared by removing the bound RNA. For the simulations of the SRSF1 RRM2 complex, we used the first frame of its NMR structure (PDB: 2M8D). (46) To increase the sampling efficiency, the simulated RNA sequence was shortened to 5’-AGGAC-3’ and we removed the unstructured N-terminal chain of the RRM2 (residues 106-119). For simulations of the A-RNA duplex, we used its X-ray structure (PDB: 1QC0; chains C and D) (48) as the start. Simulations of the free r(UUUUU) and r(UUUA) were started from the structure of the HuR RRM3 complex (44) with the protein removed. Initial structures for simulations of r(AAAA), r(CAAU), r(CCCC), and r(UUUU) tetranucleotides (TNs) and r(UCAAUC) hexanucleotide (HN) were prepared using Nucleic Acid Builder (NAB), (49) corresponding to an A-RNA duplex structure with the complementary strand removed. In all the spontaneous binding simulations (SBS), the bound RNA was manually shifted away from the protein prior to the system building so that the distance between the two molecules was ∼20 Å. We suggest this distance is entirely sufficient for randomizing RNA’s internal structure prior to its first spontaneous and random contact with the protein. Nevertheless, in case of HuR RRM3–r(UUUUU) SBS, we also used a second starting structures with different initial conformation and position of the RNA in relation to the protein.

### System building and simulation protocol

We used xLeap module of AMBER 20 (49) to prepare the coordinate and topology files. The RNA was described by the bsc0χOL3 (i.e., OL3) force field (recommended first-choice AMBER RNA *ff*). (43) For simulations of TNs and HN, the OL3 *ff* was additionally combined with modification of phosphate vdW parameters (50) and adjusted backbone dihedrals (51,52) (so called OL3CP version). For simulations of the free r(UUUUU), we also tested the Chen–García (30) and DESRES (53) RNA *ff*s. In most simulations, the protein was described using the ff12SB *ff* which is the earlier version of the ff14SB. (54) The ff12SB provides a better performance due to one specific dihedral term which significantly alters rotational kinetics of aromatic side-chains. This term is present in the ff12SB but was removed in both ff14SB and ff19SB which can cause problems in description of some protein-RNA interactions. So far, this has been only sparsely noted in the literature. (55-58) The problem is now defined and documented in this work, by comparing the performance of the ff12SB, ff14SB, and ff19SB *ff*s for the isolated HuR RRM3 protein.

Prior to simulation, all systems were surrounded in an octahedral box of either SPC/E (59) or OPC (60) water molecules with a minimal distance of 12 Å between the solute molecules and the box border. Only OPC (60) water model was used in conjunction with the ff19SB protein *ff*. (61) For TNs and HN, the rectangular water box was utilized. Excess-salt ion concentration of 0.15 M was established by randomly adding KCl ions (62) around the solute. The minimization and equilibration was performed using the pmemd.MPI module of AMBER 20 using the protocol described in Ref. (63). Afterwards, the production simulations were run using the pmemd.cuda (64) on RTX 2080ti graphic cards for a standard length of 10 μs. Multiple simulations of each system were run, with different trajectories obtained by assigning initial atomic velocities using a random seed number at the beginning of each simulation. SHAKE (65) and hydrogen mass repartitioning (66) was used in all simulations, permitting a 4 fs integration step. Long-range electrostatic interactions were described with particle mesh Ewald scheme (67) with periodic boundary conditions applied to handle the system border bias. The cut-off distance for non-bonded Lennard-Jones interactions was 9 Å. We used Langevin thermostat and Monte Carlo barostat (49). Some of the TN and HN simulations additionally included the gHBfix (26) and tHBfix (35) potentials which selectively modify stability of specific H-bond types.

### Stafix modification - rescaling the LJ interactions of RNA

After creating the topology files in xLeap, we introduced off-diagonal terms in the LJ matrix to modify selected nonbonded RNA-RNA potentials; an approach commonly referred to as nonbonded fix (NBfix). This was done automatically by an in-house python script (Supplementary Information) utilizing the parmed module of AMBER 20. In case of protein-RNA simulations, it required that unique atom types were defined for all RNA atoms which shared their LJ parameters with the protein. Afterwards, selected RNA-RNA pairwise LJ potentials (Figure 2) were modified in the topology files so that the well-depth value (ε) of their LJ potentials was reduced by a specific scaling factor. Two different scaling factors were tested. The modification is aimed at suppressing excessive non-specific RNA self-interactions such as the base-base, base-phosphate, sugar-base, and sugar-sugar stacking. As such, we refer to the modification as the “stacking fix” (abbreviated as stafix). For technical reasons, we avoided rescaling the LJ potentials related to H-bonds. See Supplementary information for full details.

**Figure 2.**
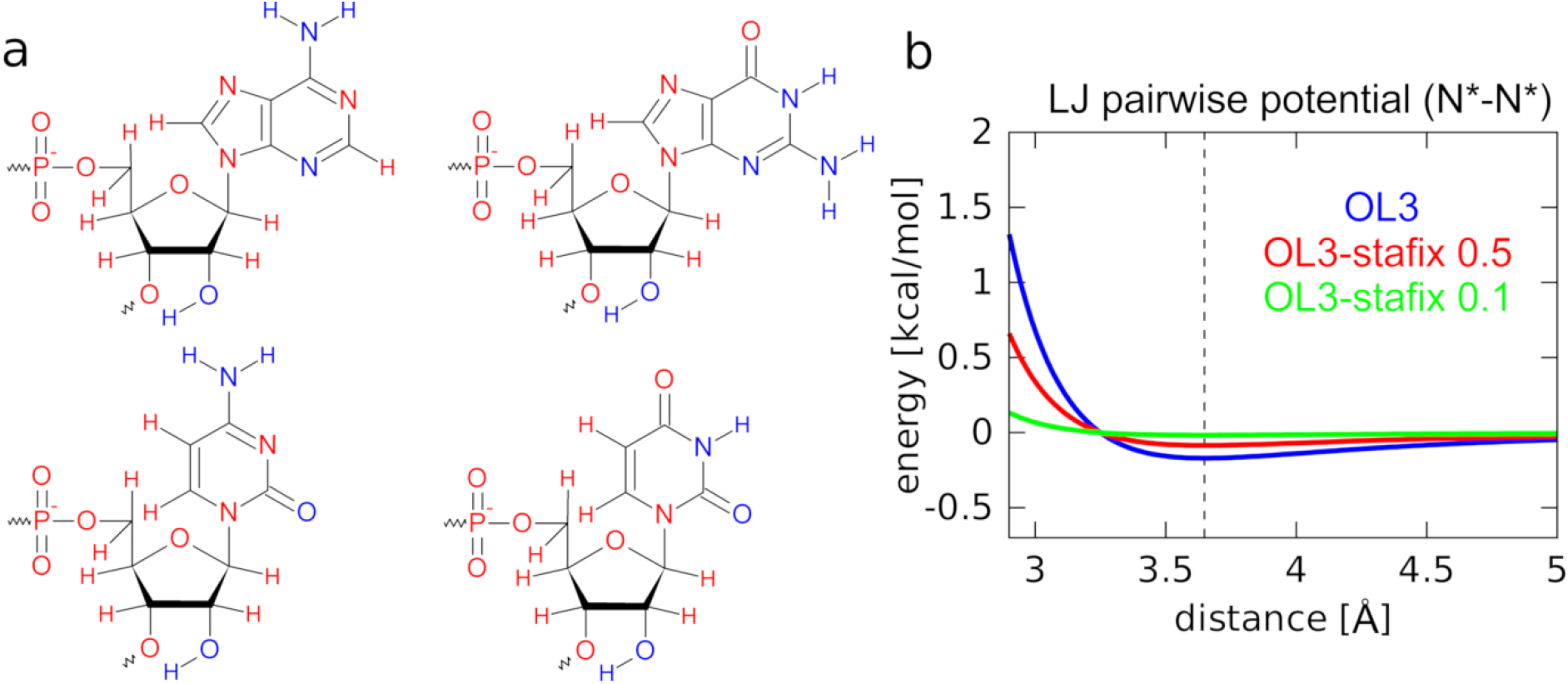
Stafix potential. a) Visualization of the standard RNA nucleotides with red and blue color referring to atoms with their LJ pairwise potentials modified and unmodified by stafix, respectively; stafix is applied between the pairs of red atoms as well as all carbons. b) Example of the unmodified AMBER LJ potential function (blue) and the same potential after applying stafix scaling factors of 0.5 (red) or 0.1 (green). Black vertical dashed line indicates the common potential energy minimum of all three functions.

### Analyses

Cpptraj module of AMBER and VMD were used for post-processing, analyses, and visual inspection of all simulations. (68,69) Povray and GNUplot were used to prepare molecular figures and graphs, respectively. For simulations of systems where we tested both the OPC and SPC/E water models, the data shown in the figures refer to systems where SPC/E was used unless stated otherwise. For graphs showing time-development data, a single simulation was selected as data source unless stated otherwise. For the histogram analyses, a combined simulation ensemble of all simulations of the given type was used and the bin size was set to 10 Å^2^. The solvent-accessible surface (SAS or SASA) of RNA and the size of protein-RNA interfaces was calculated using the LCPO method. (70) The bound protein was excluded in RNA SAS calculations. For visualization of the molecular surfaces with VMD, we used probe size of 1.4 Å.

To monitor the binding process and exchanges of bound nucleotides, we have used the following approach to identify which nucleotides occupied the specific binding pockets in simulations of HuR RRM3 and SRSF1 RRM2. We first defined list of protein heavy atoms located within 7 Å of the geometrical centers of the base aromatic rings within each pocket of the native experimental structure. The presence of individual nucleotides within the pocket in all simulation frames was then evaluated based on the distance between the geometrical center of the base aromatic ring and the geometrical centers of the previously defined lists of protein atoms. For HuR RRM3, specific nucleotide was considered to be present in the pocket when the distance was below 7 Å for pockets p1, p2, and p4 and 5.4 Å for pocket p3. For SRSF1 protein, the distance criteria was 7 Å for all three pockets. Note that this analysis of binding was based solely on empirical distance conditions and did not evaluate other criteria, such as formation of the individual protein-RNA H-bonds. Nevertheless, based on extensive visual inspection, we suggest it provides a fairly representative albeit approximate description of the binding process.

The native contacts analysis was performed by cpptraj with non-hydrogen interatomic distances in the starting structure shorter than 5 Å being considered native. For A-RNA duplex system, the base pair, base-pair step and helical parameters were analyzed by x3DNA (71) and cpptraj using default settings. MD conformational ensembles of TNs and HN were compared with available data from NMR solution experiments; (32-34) see supporting information for details. The dominant conformations of TNs and HN were identified by clustering (72) combined with the *ε*RMSD metric; (73) see also Ref. (35).

## Results

We present results of over a millisecond of standard (unbiased) MD simulations performed with AMBER RNA OL3 *ff* (43) and ones where we additionally combined OL3 with the stafix correction to eliminate the RNA over-compaction (henceforth referred to as OL3-stafix *ff*; see also Table 1 and Supplementary Table S1). The prime goal was to investigate spontaneous binding of the ssRNAs to the HuR RRM3 and SRSF1 RRM2 domains from the unbound state. The first key observation is that the RNA conformation recognized by the HuR RRM3 and SRSF1 RRM2 is not populated in simulations of the unbound RNA sequence with the OL3 *ff* due to excessive RNA self-interaction so that the ssRNA is unable to sample conformations competent for binding. As explained in the Introduction, it is an artefact typical for *ff*s parametrized to simulate folded biopolymers. This undesired behaviour was corrected with the OL3-stafix *ff* which allows routine observation of spontaneous conformational capture of the free RNAs by the proteins, atomistic details of which are for the first time described in this work. OL3-stafix also dramatically improves the stability of the protein-RNA interface when simulating the fully-bound complex. Finally, although OL3-stafix was primarily developed as goal-specific *ff* correction for protein-RNA complexes capturing unstructured RNAs, we document its effects also in simulations of TNs and A-RNA duplexes (full results are in Supplementary Information).

**Table 1.**
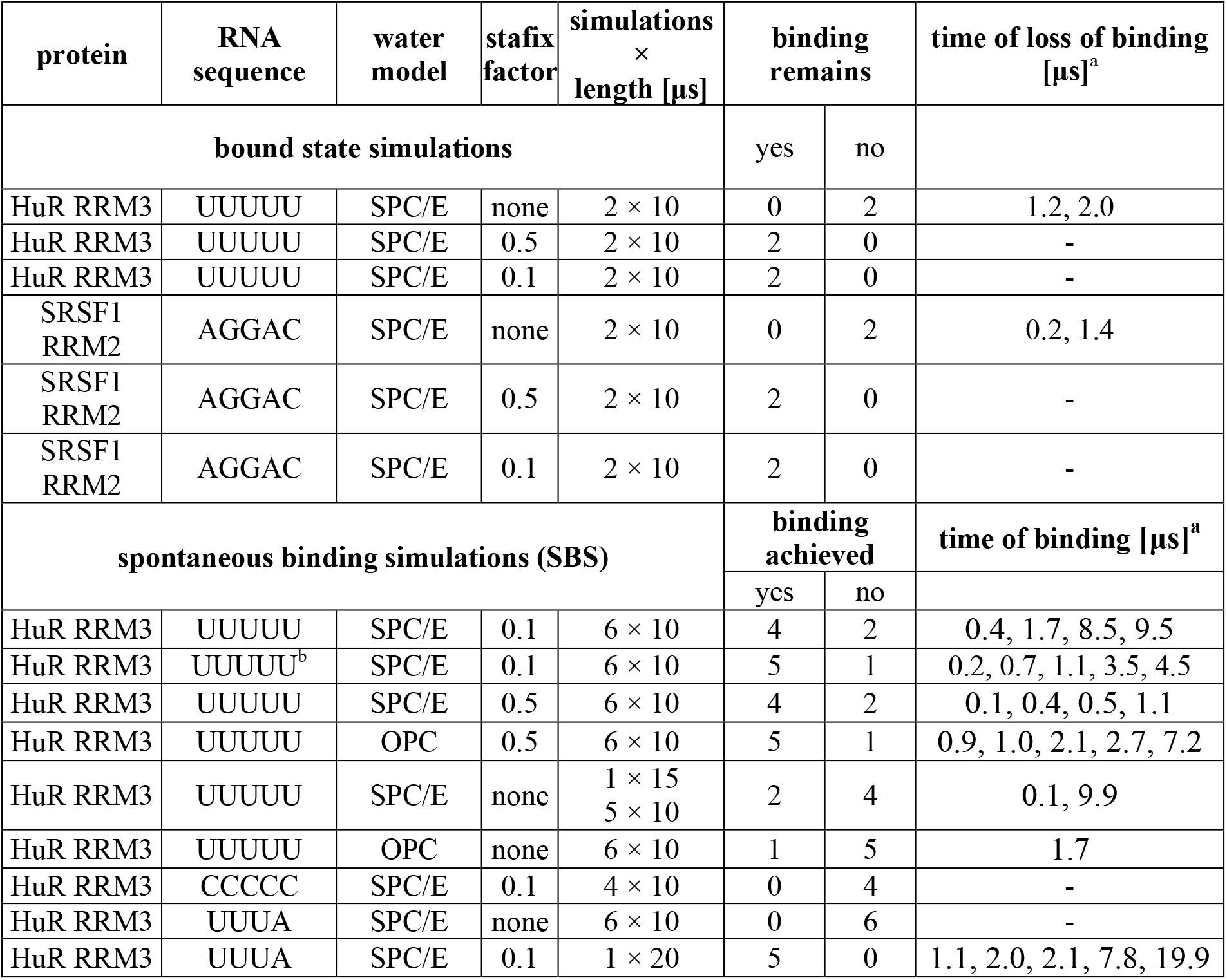
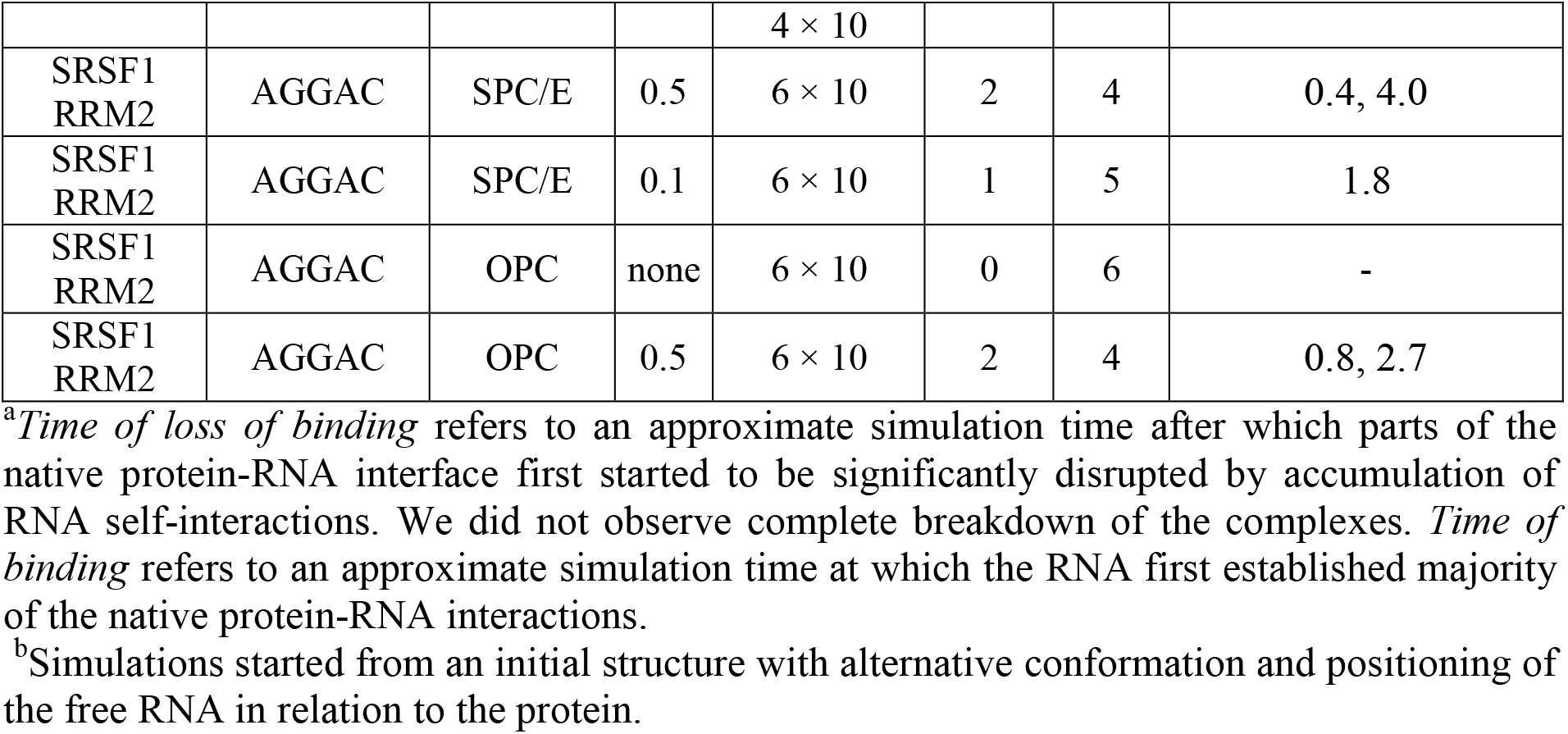
List of simulations of bound protein-RNA complexes and spontaneous binding simulations (SBS).

### OL3-stafix eliminates formation of spuriously compacted structures of free RNAs seen with the standard OL3 and some other *ff*s

MD simulations of the free 5′-UUUUU-3′ and 5′-UUUA-3′ RNAs, which are both the target sequences of HuR RRM3 (Figure 1a), showed a gradual but ultimately permanent collapse as the RNA self-interactions accumulate. We term the resulting RNA structures as ‘globules’, being best characterized by their compact globular shape (and a reduction of SAS by ∼20 % (Figure 3 and Supplementary Figure S1). This overcompaction is consistent with RNA force-field problems reported in the literature (18,22-26,32-34,53) and we observe this effect with OL3 (43) as well as DESRES (53) and García (30) RNA *ff*s (Supplementary Figure S2). Once forming the globule, the target RNAs are no longer competent to form interactions with a protein like HuR RRM3 as these require an extended RNA chain where the individual bases are accessible for readout. By applying OL3-stafix with either 0.5 or 0.1 scaling factor, we can prevent formation of the RNA globules and maintain the extended-like structure of the free RNA strand, which is accessible for the solvent, and in turn for the protein (Figure 3). OL3-stafix can also swiftly dissolve globules which had already formed (Supplementary Figure S2). We note that the OL3-stafix *ff* does not rigidify the ssRNA which we suspect could occur if we try to use dihedral reparametrizations to prevent the overcompaction.

**Figure 3.**
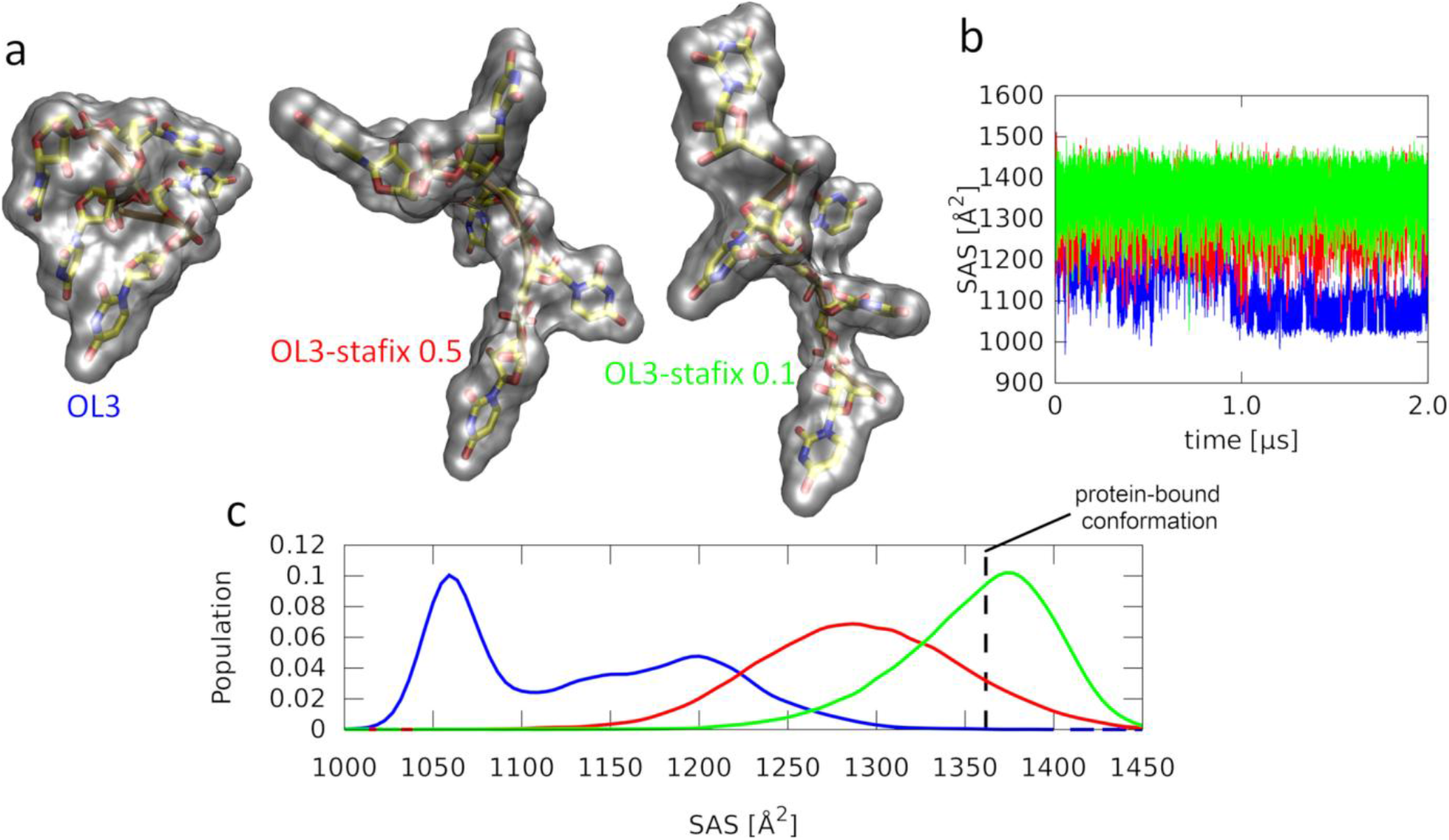
MD simulations of the free 5′-UUUUU-3′ RNA. a) Last frames of the 2-μs-long simulations (Supplementary Table S1). The transparent grey blobs indicate solvent-accessible molecular surfaces. Standard AMBER OL3 *ff* forms a compact globule (left) while more extended conformation can be maintained with both versions of the OL3-stafix *ff* (middle and right). b) Time development of the RNA solvent-accessible surface (SAS). c) Histograms of the data from panel b. The SAS value corresponding to the protein-bound conformation (vertical dashed line) is inaccessible with OL3 but regularly visited with OL3-stafix *ff*, allowing initiation of the conformational capture mechanism.

### Formation of RNA globule also disrupts bound HuR RRM3–5′-UUUUU-3′ complex

The structural collapse and SAS reduction seen for the free 5′-UUUUU-3′ RNA with the standard OL3 *ff* (Figure 3) is also affecting the RNA molecule when bound to the HuR RRM3 (Figure 4). Both experiments and MD simulations have shown that the RNA stably binds in pockets p2 and p3 while binding in pockets p1 and p4 is partially disordered (see Figure 1a and Ref. (44)). The partial-disorder makes the system prone to accumulating RNA self-interactions which subsequently degrade its interface (Figure 4a). The undesired behaviour was entirely eliminated by both OL3-stafix variants (Figure 4 and Supplementary Figure S3); see Supplementary Information for more details.

**Figure 4.**
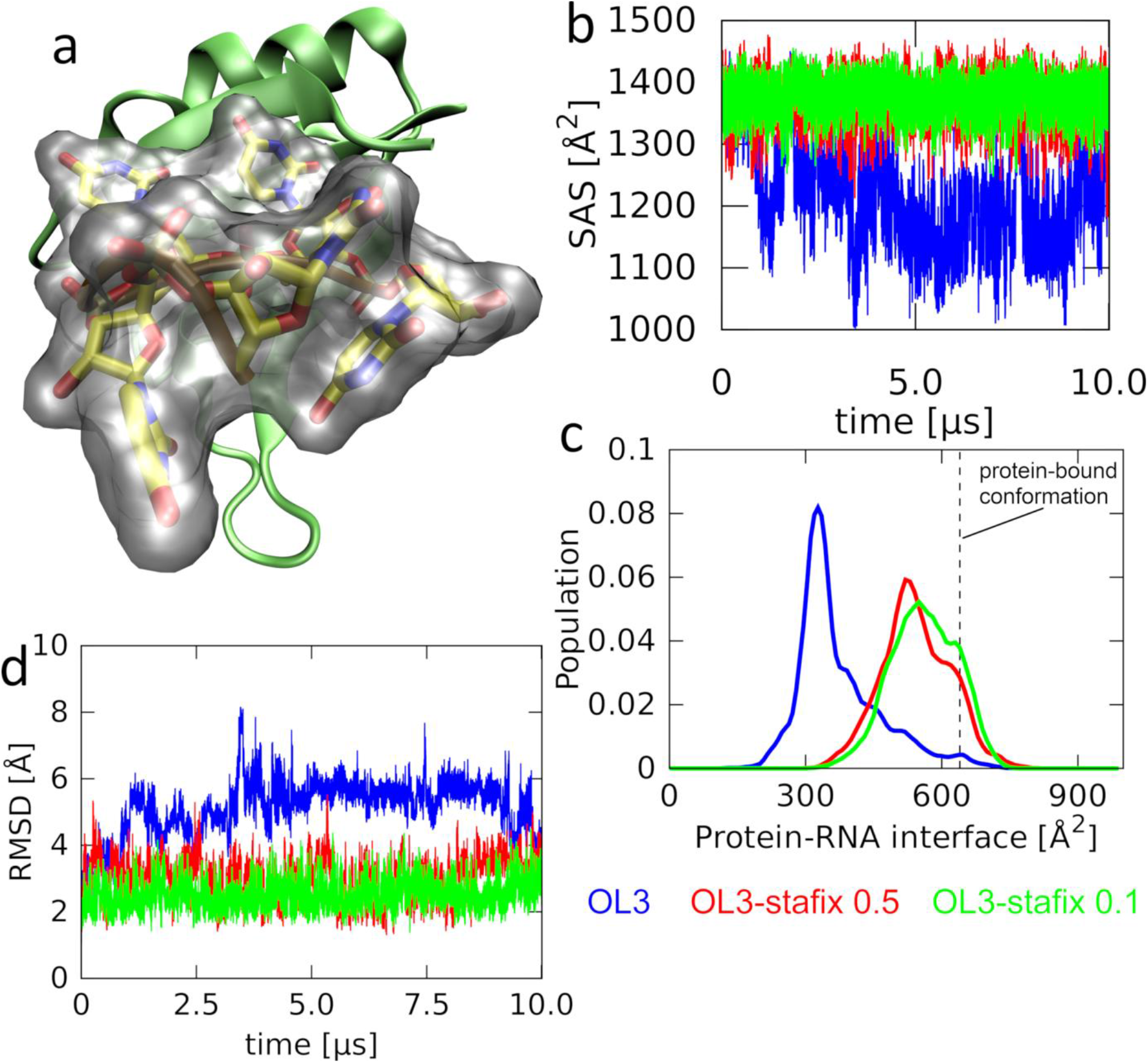
MD simulations of the HuR RRM3–5′-UUUUU-3′ complex in the bound state. a) A snapshot showing how with the standard *ff*, the interface (cf. with Figure 1a) is rather quickly disrupted in favour of spurious RNA self-interactions and a globule formation (grey blob). This is coupled with the loss of the protein-RNA interface size as it is replaced by the binding-disrupting RNA self-interactions. b) Time-development of the RNA SAS. c) Histograms of the protein-RNA interface size. d) Time-development of the RMSD for the protein-RNA interface (defined as RNA residues and protein residues within 5 Å of the RNA in the experimental structure).

### Spontaneous protein-RNA binding events of 5′-UUUUU-3′ and 5′-UUUA-3′ to HuR RRM3 on a ten-microsecond timescale

We have used the OL3-stafix *ff* in twenty-nine ten-microsecond simulations (Table 1), attempting to observe spontaneous (i.e. unguided by any pre-determined pathway or coordinate) binding events. The target RNA was placed at a ∼20 Å distance away from the protein prior to the simulation start (see Methods for details). We term such calculation as spontaneous binding simulation (SBS). RNA globules were not observed in any of the OL3-stafix SBSs (Supplementary Figure S4) and after a period of free diffusion through the water box, the RNA was successfully captured by the protein in its native binding mode in twenty-three (∼80%) simulations (Table 1 and Supplementary Figure S5).

We classify a successful binding event when most of the native binding interface is established for the rest of the simulation, leading to qualitatively identical behaviour as in simulations started from a bound structure. It includes the stable binding in pockets p2 and p3 and partially-disordered binding in pockets p1 or p4, which is also consistent with the experimental data. (44) We generally observed that once the native binding mode was established in an SBS, it remained stable (albeit very dynamical) for the rest of the simulation, demonstrating a predictive power of the stafix-modified *ff*. Also, as shown in Supplementary Information, using OL3-stafix does not abolish selectivity of HuR RRM3 to its RNA targets.

### Binding of target RNA to HuR RRM3 is a multiple-pathway multiple-step process with substantial role of structural dynamics at each step

Our SBSs of the HuR RRM3 system revealed diverse multiple pathways by which the RNA achieves native binding to the protein. Each step of the process was also fully reversible with multiple back-and-forth transitions sometimes observed. Although each binding event was unique in its details, the typical process can be characterized with four distinct steps (Figure 5). The ***first step*** involved one of the nucleotides randomly approaching and being captured by any of the four binding pockets. On a scale of tens to hundreds of nanoseconds, this was followed by the ***second step***, which was formation of what we term a ‘**pre-binding state**’, characterized by extensive but unspecific intermolecular contacts. These often involved the aromatic side-chains of the protein and the ribose sugar rings of the RNA. We suggest the pre-binding state is an intermediate between the native binding and an unbound state (Figure 5). The RNA is highly flexible in this state and samples many different interactions with the protein, including with the residues of the native binding interface. On a scale of tens of nanoseconds, this eventually triggered the ***third step*** of the binding, in which the pre-binding state transitioned into the native binding mode as multiple binding pockets became spontaneously occupied. Note that in majority of cases, this initially resulted into a misaligned binding register (non-optimal binding). For example, the U1 nucleotide would occupy the p2 pocket which allowed binding of U2 and U3 in pockets p3 and p4, respectively, thus leaving the p1 pocket unoccupied. Therefore, ***the fourth step*** of the binding process was sliding of the target RNA sequence into a binding register where all protein binding pockets are occupied. The fourth step did not always occur on the timescale of the SBS and is described in detail in the next paragraph.

**Figure 5.**
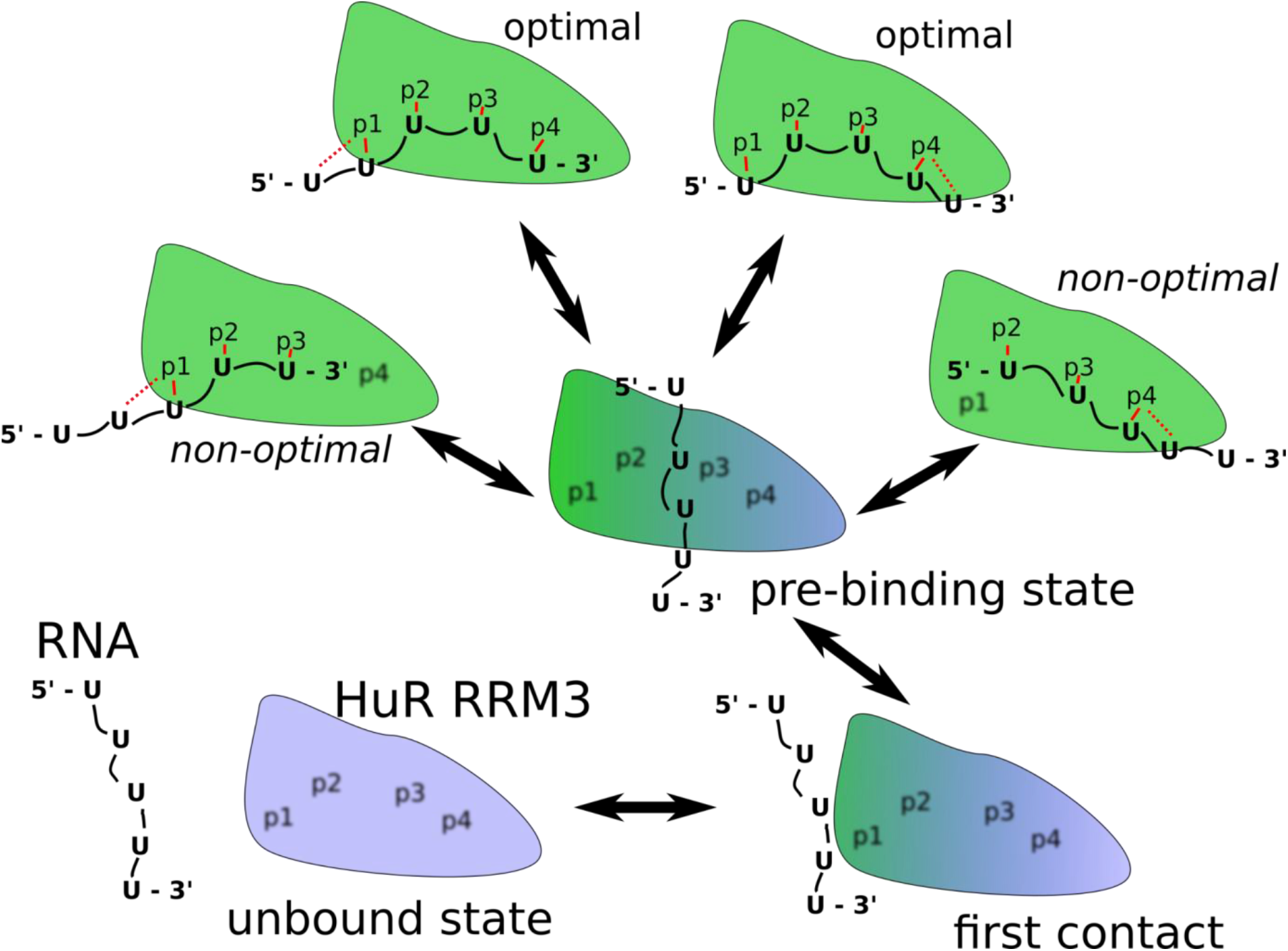
Scheme of the HuR RRM3–5′-UUUUU-3′ complex formation as revealed by SBS. The simulations start with the biomolecules separated and freely diffusing in water until one of the Us spontaneously binds in any of the pockets (first contact). This eventually brings the RNA into extensive, non-specific interaction with the protein surface surrounding the native binding pockets (a pre-binding state) which then leads to formation of the native complex interface in either non-optimal or optimal binding registers. The system can exchange between the registers by temporarily reverting into the pre-binding state. Note that the scheme is merely illustrative of the general principle and that each of the arrows actually represents multiple pathways. Similar binding pathways were followed by the 5′-UUUA-3′ RNA which, however, has only one optimal and one non-optimal binding register. The solid red lines indicate nucleotide interaction with the pocket. Dashed red lines indicate local competition between two nucleotides for a single binding pocket. Blurred pocket labels indicate empty binding pockets. Blue and green colors of the blob indicate unbound and bound protein structures, respectively.

The above mentioned four steps roughly delineate the most common scenario of the binding events observed for the 5′-UUUUU-3′ or 5′-UUUA-3′ RNAs and also allow us to estimate the associated timescales. Namely, the first step was the most limiting as without any guiding force, the RNA would sometimes spent microseconds exploring various non-native interactions with other parts of the HuR RRM3 surface or diffusing through the water box before first approaching the native binding pockets. In some of the SBSs, the first step did not occur on the ten-microsecond timescale. It was the most common cause of failed SBSs, i.e. trajectories where no successful binding events were observed (Supplementary Figure S5, #6 #12). Even when the first step occurred, another limitation was that the RNA could be captured in a position unsuitable for forming the rest of the native interface, such as the reversed 5′–3′ polarity of the RNA chain or the U1 nucleotide being captured in pocket p4 (not shown in Figure 5). In most such cases, the RNA would still eventually transition into the pre-binding state (second step) and proceed on with the binding process. However, sometimes the RNA would not depart from these unproductive and rather off-pathway arrangements for the rest of the SBS (Supplementary Figure S5, #5 #18 #51).

Trajectories where the first nucleotide was captured by pocket p4 were the most sensitive to this behaviour as pocket p4 is relatively unselective and can accommodate nucleotides in multiple, even non-native, spatial orientations. This problem may have been exaggerated by suboptimal force-field parameters of the sulphur atom of cysteine 245 which is part of the pocket p4 (Supplementary Information). Lastly, the sliding of the RNA into the full (optimal) binding register often did not occur on the trajectory’s timescale and only part, albeit a majority, of the native interface remained occupied until the end in most SBSs (Supplementary Figure S5, #1 #8 #48). We still considered SBS with such non-optimal binding interfaces as successful. As explained below, we actually suggest that the native bound state of the 5′-UUUUU-3′ consists of two optimal and two non-optimal binding registers (Figure 5).

### Structural basis of the 5′-UUUUU-3′ RNA sliding within the binding register of HuR RRM3

In principle, the individual Us of 5′-UUUUU-3′ RNA can equally-well occupy any of the four binding pockets of HuR RRM3 (Figure 1a) which raises a question how the optimal binding register is established. Indeed, our SBSs showed that the initial binding events often result into non-optimal binding registers. Most commonly, three pockets were filled while one (p1 or p4) remained empty (Figure 5, Figure 6a and Supplementary Figure S5) and in many SBSs, this was the final state of the binding process observable on the ten-microsecond timescale. However, there were several instances where the RNA later spontaneously shifted within its binding register to reach the optimal state with all four pockets filled (Supplementary Figure S5, #7 #9 #13 #22 #47). We refer to this process as sliding.

**Figure 6.**
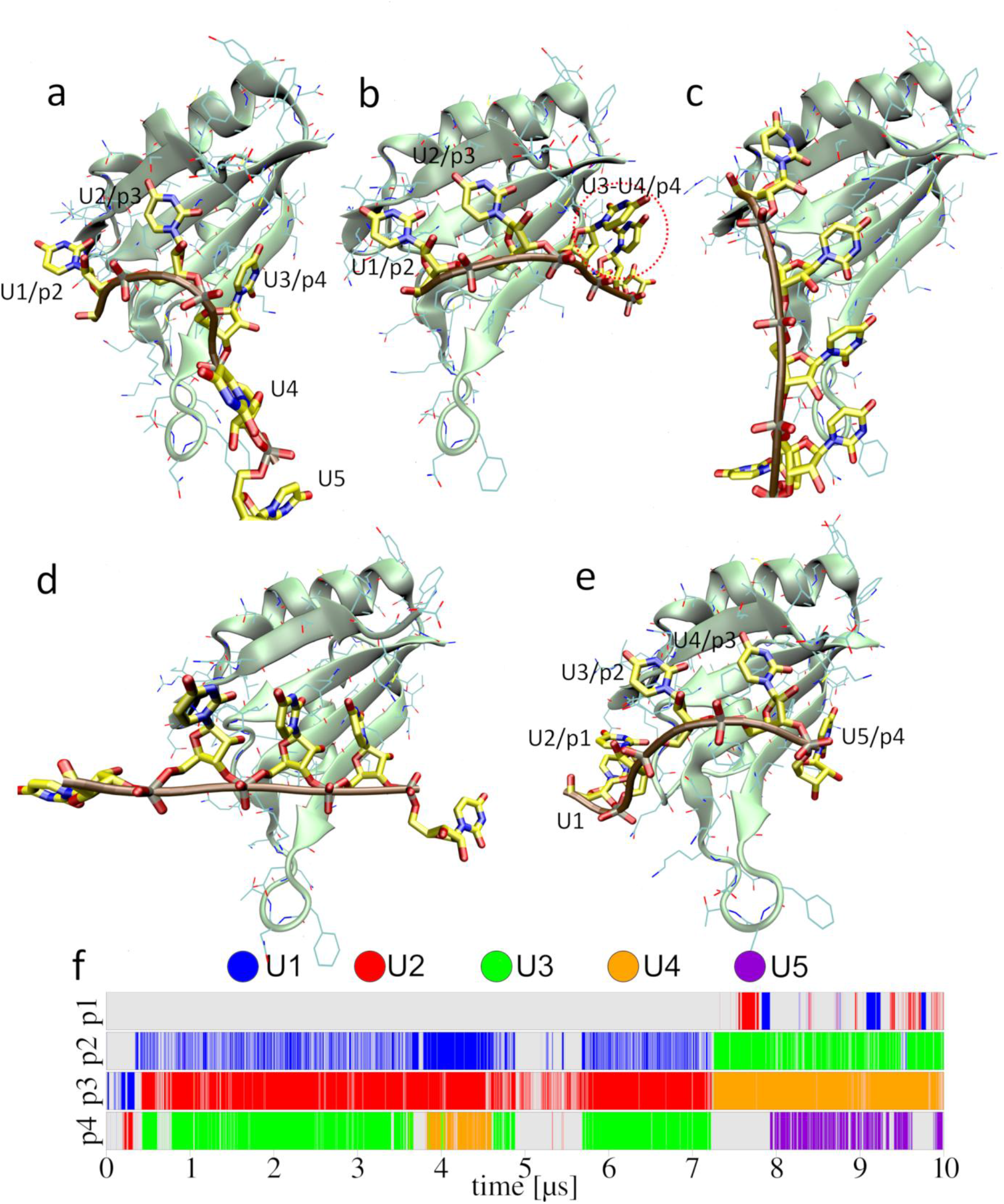
Spontaneous 5′-UUUUU-3′ RNA sliding on the surface of HuR RRM3 in one simulation. After first initial contact with the protein at ∼0.24 μs, the RNA bound in a non-optimal binding register with only three out of the four pockets filled (a). It was followed by many reversible local disruptions/fluctuations of the protein-RNA interface where U3 and U4 competed for the p4 binding pocket (b; red circle). This eventually led to a larger-scale global disruption at ∼4.9 μs where the RNA detached from the binding pockets but did not fully depart the protein’s surface, entering the pre-binding state. The system would repeatedly fluctuate between the pre-binding and native states until ∼7.1 μs when one of the global disruptions was followed by a sudden chain sliding by two nucleotides (d; ∼7.25 μs) in the 5′ direction and the native interface was (re)formed with all four binding pockets regularly occupied though still dynamically fluctuating (e). f) Time-development of the binding pocket occupancies by specific nucleotides; note that the binding-occupancy descriptor does not fully visualize the dynamicity of the complex.

Similarly to the binding, the individual sliding events also proceeded in a unique fashion, indicating it is a very multidimensional multiple-pathway process which does not appear to be easily describable by some collective variables, simplified coordinates or descriptors without an information loss. Nevertheless, we can say that the sliding (register exchange) is preceded by a local competition between two nucleotides for a single binding pocket (Figure 6b and Figure 5), most commonly the pockets p1 or p4 which are partially-disordered and able to transiently accommodate two nucleotides simultaneously. This local competition can trigger a large perturbation of the native interface in which the system temporarily transitions back into the pre-binding state (Figure 6c). As previously mentioned, the RNA is very flexible while in the pre-binding state. It tends to quickly revert back towards the native binding state but in some cases, it does so with a binding register shifted by one or more nucleotides (Figure 6d), thus completing the sliding process (Figure 6e). The register-exchange dynamics appears to be a process regularly occurring on 1-10+ µs timescale.

### Spontaneous binding of the SRSF1 RRM2–5′-AGGAC-3′ complex

Unlike the partially-disordered interface in the HuR RRM3 system, the SRSF1 RRM2 recognizes GGA triplet motif in mRNA via classical well-defined binding pockets. (46) There is also no viable alternative binding register as the G_2_, G_3_, and A_4_ nucleotides of 5′-AGGAC-3′ RNA can only fit into pockets p1, p2, and p3, respectively (Figure 1b). We observed successful binding in five out of eighteen SBSs (Table 1 and Supplementary Figure S6); a noticeably lower success rate (∼30%) than for the HuR RRM3 system. This was mainly due to the abundance of non-native sites explored by the RNA on the SRSF1 RRM2 surface which sometimes prevented it from reaching the native binding interface on the 10-μs-timescale. In other words, the binding landscape of the SRSF1 RRM2 - 5′-AGGAC-3′ complex appears to be considerably frustrated by off-pathway binding. The second reason was that there is no non-optimal binding register possible for the 5′-AGGAC-3′ RNA and it needs to achieve the optimal binding register in context of a single binding event instead of being able to reach it incrementally via sub-optimal binding registers like in the HuR RRM3. Likewise, there was no indication of any pre-binding state (Figure 5) which could guide the RNA towards the native binding interface. Indeed, in the five successful SBSs, we always observed one of the nucleotides spontaneously binding in one of the native binding pockets which then lead to binding of the second and finally the third (Figure 7). Although relatively straightforward compared to HuR RRM3 (see above and Figure 5), the RNA binding to SRSF1 RRM2 is still a multiple-pathway process as it does not appear to have any preference for order in which the nucleotides are bound. The leading cause for failure among the unsuccessful SBS was the RNA not reaching the native binding site within the simulation timescale (Supplementary Figure S6, #4 #12). Another common reason was one of the nucleotides binding in compatible but non-native binding pockets (i.e. A_1_ in p3 or G_3_ in p1; Supplementary Figure S6, #5 #21). Such binding was relatively stable on the simulation timescale and stalled the RNA, preventing it from reaching the native binding mode. Importantly, there were no RNA globules observed in SBSs when using OL3-stafix (Supplementary Figure S7) and we also observed improved performance of simulations started from a fully bound complex (Supplementary Figure S8); see Supporting Information for details.

**Figure 7.**
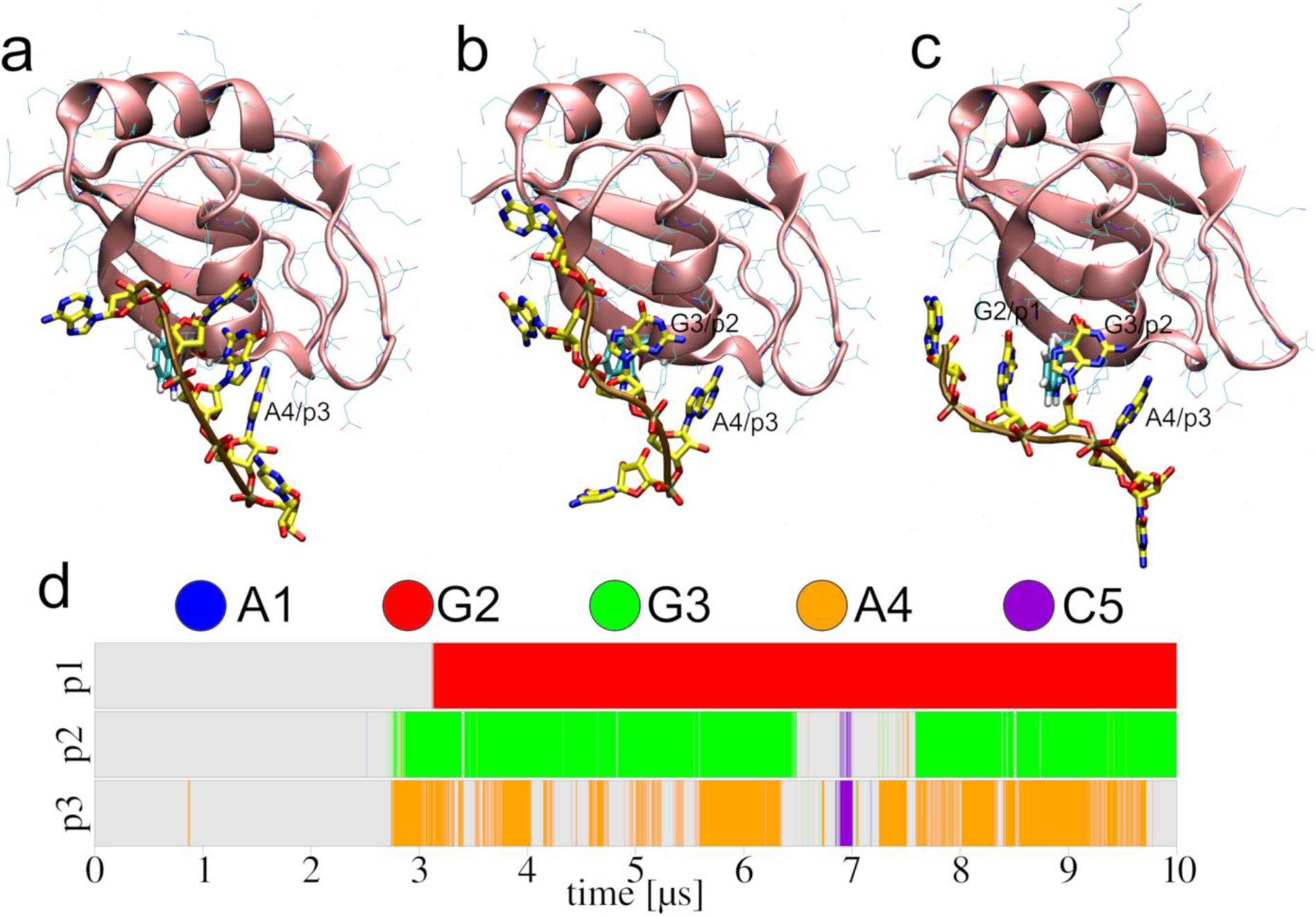
Spontaneous binding of 5′-AGGAC-3′ to SRSF RRM2 as observed in one of the OL3-stafix SBS. The A_4_ nucleotide bound in the third pocket (a; ∼2.70 μs), followed by G_3_ in the second pocket (b; ∼2.87 μs) and finally G_2_ in the first pocket (c; ∼3.13 μs). d) Time-development of the binding pocket occupancies by specific nucleotides.

### Standard *ff* does not allow binding

For comparison, we also performed SBSs where we did not use stafix, i.e. with the standard OL3 *ff*. Although the RNA would sometimes sample the native binding pockets, successful binding occurred only in three SBSs out of eighteen and only for the HuR RRM3–5′-UUUUU-3′ system. The successful binding can rarely occur even for the RNA globule when some uracils become temporarily exposed on its surface while simultaneously being at the proximity of the native binding site. However, even then the binding is visibly frustrated compared to simulations with OL3-stafix due to competition from the RNA self-interactions which ultimately overpower the binding tendency and the binding is lost again. In all other simulations, including the SRSF1 RRM2–5′-AGGAC-3′ or HuR RRM3–5′-UUUA-3′ systems, extensive formation of RNA globules entirely prevented stable binding with standard OL3 *ff*. Full details are in the Supplementary Information.

### Influence of water model and selection of protein *ff*

We observed virtually identical performance when using the SPC/E (59) or OPC (60) water models in our simulations (Table 1 and Supplementary Table S1), including the tendency to form RNA globules without stafix (Supplementary Figure S9) and the similar benefit provided by the modification. Therefore, we suggest that both water models are equally valid choices for SBS. In all our simulations, we used the ff12SB *ff* to describe the proteins. The reason for preferring this protein *ff* version over the newer ff14SB or ff19SB is the superior (for the present systems) description of the phenylalanine and tyrosine side-chains provided by ff12SB. These residues are prominent at the protein/RNA interfaces of RRM domains studied in this work (44,46) as well as protein-RNA complexes in general. Full details are in the Supporting Information.

### Simulations of tetranucleotides, hexanucleotide and A-RNA

OL3-stafix is a goal-specific *ff* modification in no way intended to be a multi-purpose RNA *ff*. Still, we also examined its performance in simulations of tetranucleotides (TNs), hexanucleotide (HN) and A-RNA duplex (Supplementary Table S1) – simple RNA motifs which are commonly used in force-field benchmark studies. As expected, the use of OL3-stafix increased the RNA SAS of all simulated TNs and the HN. On the other hand, simulations of the A-RNA duplex revealed only minimal changes. Full details are in the Supporting Information.

## Discussion

### Spontaneous binding simulations (SBS) of protein-RNA complexes: from two-state model to ensemble description

We have used MD simulations to explore completely spontaneous protein-RNA binding events for two simple protein-RNA interfaces (Table 1). Such calculations were previously accomplished for other systems by either using coarse-grained models or by molecular docking approaches (10,18,74) but not in the context of unbiased full-atomistic MD. The simulations reveal astonishingly rich and diverse multidimensional dynamics on timescales from nanoseconds to many microseconds. To achieve this, we developed a force-field modification (called stafix) which addresses the tendency of unstructured ssRNAs to excessively self-interact in MD simulations when using AMBER *ff*s designed for folded RNA structures. Using this method, we were for the first time able to observe the entire binding process of ssRNAs to HuR RRM3 and SRSF1 RRM2 (44,46) proteins, from the free diffusion around the proteins through initial conformational capture to final formation of the native binding interfaces by induced fitting.

### Binding of RNA to HuR RRM3 is a complex multiple pathway process with partially disordered bound state

It can be said without exaggeration that no two binding events that we observed for the HuR RRM3 system proceeded in exactly the same fashion. Each binding event was unique. This demonstrates the value of standard (unbiased) MD simulations. To describe all nuances of the binding process using a limited number of collective variables would be an arduous task and the dimensionality reduction excessive. (75) This was most obvious with the 5′-UUUUU-3′ RNA which can bind to HuR RRM3 in multiple binding registers while the individual Us can in principle occupy all four binding pockets equally well. As a consequence, there are countless pathways between the unbound and bound states that the system can utilize during the binding process. Importantly, the multidimensionality is also a hallmark of the bound state. The bound complex is characterized by rich multidimensional thermal dynamics involving four registers and ranging from subnanosecond to (multi)microseconds time scales. The RNA is localised at the binding site but semi-disordered.

In our HuR RRM3 SBSs, the binding typically started with one nucleotide randomly binding in one of the pockets which brought the rest of the RNA close enough to also sample the other pockets and eventually bind there. At this stage, we also could observe an intermediate “pre-binding state” where the native binding pockets are not yet filled but the RNA is already in extensive molecular contact with the protein surface proximal to the native binding interface.

The initial binding events almost always resulted in non-optimal binding register where not all the binding pockets were filled (Figure 5). In few simulations, we later saw the RNA spontaneously shifting into the optimal binding register, i.e. the sliding (Figure 6). The sliding events were precipitated by the RNA temporarily transitioning back into the pre-binding state before returning into the native binding state with an exchanged register (Figure 6). We suggest the HuR RRM3 might be able to rapidly scan the target RNA sequence in this fashion. Firstly, the RNA pre-binding state (Figure 5), in which the RNA is very flexible compared to the native binding state, allows sliding without the need to fully separate the RNA from the proteins surface. This is advantageous as full separation of the two biomolecules would dramatically decrease solvent entropy (76,77). The pre-binding state is evidently separated only by relatively small free-energy barrier from the native binding state, allowing it to be regularly visited on a microsecond timescale. It thus can be considered as a genuine part of the binding funnel indistinguishable from the bound state at lower time-resolutions. The rich dynamics is greatly aided by the partial binding disorder of pockets p1 and p4, which facilitates competition between neighbouring nucleotides and results in a very flat multiple-minima and multidimensional free-energy surface of the bound state (Figure 6).

Secondly, when the sliding occurs with a single concerted movement in the context of the diffusive pre-binding state, HuR RRM3 avoids the speed limitations of having it proceed one nucleotide at the time. Although such process of sliding “pocket after pocket” is in principle possible, a complete sliding event would require all five nucleotides exchanging in the same direction (either 5′ to 3′ or 3′ to 5′). This “linear diffusion” would be controlled (hindered) by multiple barriers, leading to a quite slow exchange rate unusable in biological context; it is often referred to as the speed-stability paradox of the 1D diffusion (Figure 8a). (78) It is the presence of pre-binding state that allows the sliding.

**Figure 8.**
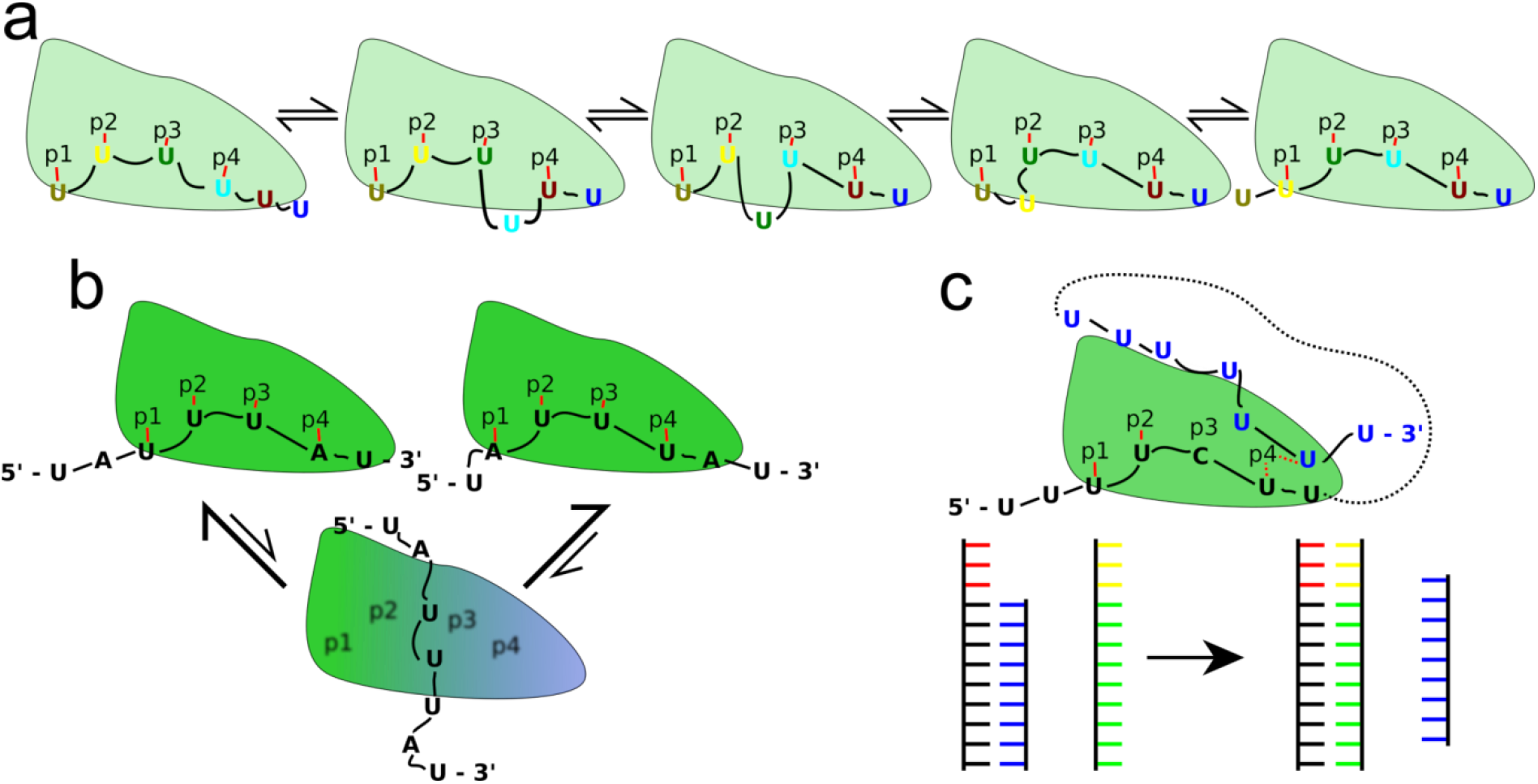
Possible mechanisms of the RNA movements and binding register exchanges in HuR RRM3 system. a) One-dimensional “inchworm” sliding movement of the poly-U hexamer RNA. Although very intuitive, such model is unlikely to have biological significance due to very slow translocation speed arising from reversibility of its individual steps, presence of multiple barriers, and lack of the external driving force. Our simulations indicate that instead of the inchworm model, the HuR RRM3 undergoes reversible transitions to the pre-binding state to facilitate an unobstructed barrierless sliding or diffusion (Figure 5 and Figure 6). b) Binding of the ideal 5′-AUUUA-3′ motif to HuR RRM3 and hypothetical exchanges between its two possible binding registers with the pre-binding state as the intermediate. c) Strand displacement of the non-target (bound) RNA (black) by the target (incoming) RNA (blue). The target RNA can be either a separate molecule or a sequence further downstream or upstream of the non-target sequence separated by a linker of unspecified length (black dashes). The partially-disordered p4 pocket can temporarily accommodate two nucleotides and could serve as a bridgehead for the incoming RNA, thus greatly accelerating displacement of the bound sequences. The ladder-like model below shows the analogous toehold scenario of strand exchange in DNA and RNA duplexes. (82-84) The single-stranded segment of the duplex (i.e. the toehold; red) allows the invading strand (green and yellow) to rapidly displace the original strand (blue). The partially-disordered pockets p1 and p4 in HuR RRM3 could function similarly to the duplex toeholds and accelerate the RNA exchange.

Thirdly, a two state model of the bound state with the native state (where sliding is slow) and the pre-binding state (where sliding is fast) means that the protein may effectively separate affinity (or binding capability) from specificity and is able to separately tune both during evolution. The essence of the sliding mechanism employed by HuR RRM3, i.e., local disruption followed by quick global disruption followed by the register shift and restoration, resembles an earlier described translocation mechanism of the WRKY domain protein along DNA helix. (79) We suggest it could be a general strategy employed by proteins specialized on efficient scanning of long nucleic acid sequences without external driving force. In case of HuR RRM3, it would allow it to rapidly search for the RNA sequence whose binding corresponds to thermodynamic minimum while efficiently screening against utterly incompatible sequences (such as the poly-C RNA, see the Supplementary data) by sequestering them in the non-specific and relatively weak pre-binding state.

In summary, binding of ssRNA to HuR RRM3 is a textbook example of a process whose full understanding requires the ensemble description. (80,81) It should be considered in MD simulation studies as well as in interpretations of experiments. (44,45)

### Binding to HuR RRM3 in context of multiple or full-length mRNAs

Similarly to most *in vitro* experimental studies, our present work does not consider how competition between multiple RNAs for the same binding site would impact the suggested binding mechanism (Figure 5). We aim to be able to perform such calculations in the future but additional *ff* modifications may be required to make them feasible which is already outside the scope of this paper. Nevertheless, the present data along with earlier published experimental studies (44,45) allow us to formulate several hypothesis.

Firstly, it was shown that the ideal target sequence for HuR RRM3 is 5′-AUUUA- 3′. (44) This sequence can bind in two binding registers (with either A_1_ or A_5_ nucleotide in pockets p1 and p4, respectively) which likely provides entropic advantage. Based on our simulations, we suggest that the pre-binding state (Figure 5) might represent a low barrier pathway between the two binding registers (Figure 8b), giving rise to a very broad free-energy minimum when binding the 5′-AUUUA-3′ and stalling the conformational search at this sequence.

Secondly, we suggest that besides a linear scanning, a strand replacement mechanism scenario may play a role, in which a bound RNA would be displaced by a second RNA or by sequence upstream or downstream of the bound nucleotides (Figure 8c). Although this process cannot be simulated at the moment, we suggest that the observed partially-disordered nature of pockets p1 and p4 and their ability to temporarily accommodate two nucleotides (Figure 5 and Figure 6) would facilitate such a scenario. By providing an inherent bridgehead for the incoming RNA, the speed of the exchange in favour of a strand with thermodynamically more stable binding would be accelerated rapidly, akin to the toehold or end fraying mechanisms well-known for strand displacements in DNA and RNA duplexes (Figure 8c) (82-84) or the active cycling models of RNA exchange on the surface of the bacterial Hfq protein. (57,85,86)

### Binding strategy of SRSF1 RRM2 is very different from HuR RRM3

Our SBSs identified no pre-binding state for the SRSF1 RRM2 system. As a consequence, the SBS success rate was lower for this system (Table 1) as the conformational capture and the induced fit needed to form the native interface had to occur in essentially a single uninterrupted “all or none” process (Figure 7), which presents a sampling limitation. Since there were no regular sliding events observed, the correct binding register also had to be established on the first try. For example, we could observe a situation where the second G of the GGA triplet bound in the first pocket. Without sliding, the only way to resolve such situation was full separation and another binding attempt which was very slow and often did not occur on the ten microsecond timescale of our simulation. We cannot rule out that this behaviour could be a consequence of using a very short RNA which contains only a single GGA triplet. In full mRNA, the competition from the other nucleotides might be used to dislodge the out-of-the-register binding (see, e.g. Figure 8c). However, we suggest that our simulations might reflect genuinely different evolutionary requirements placed on the SRSF1 RRM2 and HuR RRM3 proteins. In any case, the simulations suggest that different proteins may use diverse mechanisms and pathways even for the at first sight straightforward ssRNA binding, indicating that there is no universal mechanism by which RRMs search for their RNAs.

### Stafix potential prevents formation of RNA globules

The inability to simultaneously describe folded and disordered (unstructured) biopolymers is an Achilles’ heel of contemporary biomolecular *ff*s. (21) In this work, it is showcased by the ssRNA collapsing into hyperstable globules which outcompete protein-RNA interactions, making such ssRNA unsuitable for protein-RNA binding (Figure 3). The RNA seemingly prefers self-interactions over any other form of intermolecular contact so that the RNA self-interactions gradually disrupt even initially bound complexes (Figure 4 and Supplementary Figures S3 and S8). In other words, the global *ff* thermodynamic minimum likely corresponds to the RNA self-interactions. We addressed this problem with a force-field modification altering the stability of RNA-RNA vdW interactions via rescaling selected pairwise LJ potentials (Figure 2); technically using an approach which is known as nonbonded fix (NBfix). (87) NBfix is very effective for separate tuning of interactions which cannot be optimally described using a single set of LJ parameters. (30,38,88-92) We call the new modification *stafix* (<u>sta</u>cking <u>fix</u>) as it primarily addresses the overstabilization of stacking present in the standard OL3 *ff*. Simulations with OL3-stafix can be used to observe spontaneous binding as well as to stabilize protein-RNA interfaces (Table 1 and Supplementary Figures S5 and S6). However, it could also be of interest in simulations of free ssRNAs (Supplementary Information).

The suggested design of stafix (Figure 2) is a result of multiple iterative trial-and-error attempts. Our original intent was to only weaken the excessive base-base stacking which we suspected was the chief factor in formation of the spurious RNA self-interactions. (18,93) This initial approach proved successful in lowering the amount of base stacking, however, it immediately unmasked similarly excessive sugar-sugar, sugar-base, as well as base-phosphate stacking and vdW interactions; an observation consistent with the earlier works. (38,90,92) Thus we ultimately extended the list of rescaled atoms to weaken stacking (or vdW) interactions of the entire RNA. Note that we do not suggest OL3-stafix as a general *ff* but rather a task-oriented modification focused on achieving two specific goals. Firstly, it represses RNA globules and provides ssRNA ensemble populations of the extended unstructured conformation which is commonly recognized by the proteins but rarely populated with the standard *ff*. Secondly, it speeds up RNA’s conformational transitions by smoothing its folding landscape. It actually could be a biologically relevant description for the short ssRNAs, (94) rather than the likely excessively rugged folding landscape predicted by the standard *ff*. (18,95) Fine-tuning the individual details in design of stafix (Figure 2 and Supplementary Information) is relatively unimportant as long as the overall cumulative effect is achieved with minimum or no side-effects for a given set of goals (we reiterate that our goal was not to create a universally applicable core RNA *ff*). We obviously do not recommend the use of OL3-stafix for structured RNAs. In fact, our preliminary tests (data not shown) on the Neomycin-sensing riboswitch which contains a highly-conserved U-turn motif (96) indicated a suboptimal performance with OL3-stafix. However, it is also encouraging that for the A-RNA duplex we observed only very minor changes of helical parameters (Supplementary Table S3). We suggest that stafix rescaling factor 0.5 (Table 1) should be sufficient to eliminate the ssRNA globules, allowing many simulation studies that are right now not doable with the standard *ff*.

We also showed that OL3-stafix very efficiently eliminates the so called intercalated conformations of the TNs, suggesting that this spurious conformation at least partly originates from the force-field terms addressed by stafix. At more modest scaling factors, stafix could be of some interest for future force-field development, in particular when combined with the other methods proposed to improve simulations of ssRNAs, such as the gHBfix. (26,36) However, its greatest merit is without any doubt with simulations of protein-RNA complexes where the unstructured ssRNA often corresponds to the RNA conformation recognized the protein. We suggest that elimination of the compacted RNA structures from the free-energy landscape in such studies is justified. Even if such non-specific self-interacting RNA structures are to certain extent populated in the ensembles of real molecules, such RNA states are not likely to contribute to productive binding pathways, assuming the conformational capture mechanism at the initial stages of the binding. In other words, they are localized off-pathway from the binding funnel on the free-energy landscape.

In the present study, the stafix modification was used with the OL3 AMBER *ff* (termed as OL3-stafix), but in principle it could be combined also with other nucleic acids *ff*s, including those for DNA. Considering the known difficulties to reparametrize the general RNA *ff*, (18,26,36) extending established core multi-purpose *ffs* such as OL3 by goal-specific modifications may allow to carry out many insightful simulation studies which are not otherwise executable. This is a common strategy in coarse-grained modeling that could be used also in the framework of atomistic *ff*s.

### Future challenges and perspective

Many of our SBS trajectories revealed various non-native but long-lived protein-RNA interactions sampled by the system which prevented it from reaching the native binding. This was quite visible with the SRSF1 RRM2 system (Table 1). We suspect that nonspecific intermolecular interactions, albeit to a lesser degree, might be similarly excessive as the RNA self-interactions. In fact, this is to be expected as the vdW parameters for proteins and RNA are largely identical. Therefore, studies of more complex protein-RNA interfaces may require additional goal-specific adjustments of the methodology to destabilize off-pathway non-specific binding sites; the presently simulated systems represent relatively simple protein-RNA interfaces. Also, use of enhanced-sampling methods may become inevitable to better focus the SBS on the binding funnel in more complicated systems although extreme care will have to be taken not to excessively reduce dimensionality of the binding process. In any case, it is evident that atomistic MD simulations can provide striking insights into the ensemble nature of protein-RNA binding which is difficult to assess by other techniques.

## Conclusions

In this work, we for the first time sampled a full binding funnel of two protein-RNA complexes – HuR RRM3 and SRSF1 RRM2, using completely unbiased full atomistic MD simulations. We provide structural details of the protein-RNA complex formation and binding register exchanges. The simulations furnished the first insights into complexity and diversity of dynamical processes that can occur at protein-RNA interfaces. They underscore the importance of ensemble-based description of protein-RNA interactions, which especially for the HuR RRM3 complex is essential to understand not only the binding process, but also properties of the partially disordered bound state. On methodology side, our work required development of a force-field modification called stafix which eliminates RNA’s tendency to spuriously self-interact. Stafix provides long-term stabilization of protein-RNA interfaces and allows simulations of the binding process. After eliminating the spurious RNA self-interactions, the simple non-polarizable *ff* becomes a surprisingly powerful tool for prediction of protein-RNA interactions and complex formation mechanisms, at least for some interfaces.

## Supporting information

Script

## Supporting Information

Additional details on simulation methods and results. Supplementary Figures and Tables. Python script for implementation of stafix in AMBER MD simulations.

## Data availability

The raw MD simulation trajectory data is available from the corresponding author upon request due to large size of the datasets.

## Acknowledgement

We acknowledge the use of CESNET data storage facilities [grant number LM2018140].

