## Supplementary for "Spontaneous binding of single-stranded RNAs to RRM proteins visualised by unbiased atomistic simulations with rescaled RNA force field"

### Supplementary Text

**Design of the stafix potential.**

When designing the stafix potential, we purposefully excluded the H-bond donor groups from the list of interactions rescaled via NBfix (main text Figure 2), in order to not interfere with H-bonding. Exclusion of H-bond acceptors would have the same desired effect but the far larger number of such groups in nucleic acids made it the less convenient choice.

Important reason to avoid modification of H-bonding with stafix was that lowering the well-depth "flattens" the LJ function in both attractive and repulsive regions (main text Figure 2b). This leads to weakening of the non-specific vdW interactions, where the LJ potentials always facilitate attraction. However, the attractive part of H-bonding interaction is mainly facilitated by favorable Coulombic interaction between atoms with large and opposite partial charges (i.e. H-bond donors and acceptors) while LJ potential provides repulsion. Therefore, applying stafix for H-bonds can cause these interactions to be overwhelmed with the attractive Coulombic interaction which is insufficiently compensated with LJ repulsion, leading to unnaturally short interatomic H-bond distances and actual overstabilization of H-bonds rather than their weakening. Obviously, this would further exacerbate the problem with spurious RNA self-interactions which is why we did not include H-bond interactions into the stafix. Note that to modify stability of H-bonds one can use, for example, the gHbfix potential. (1,2) The gHbfix can be easily applied alongside stafix. Lastly, we did not apply stafix to base atoms whose repulsion with the sugar contributes to the *syn*/*anti* balance as the initial tests revealed overpopulation of the *syn* state in such simulations. Thus, we excluded O2 and N3 atoms for pyrimidine and purine nucleotides, respectively. In principle, this problem could also be solved by modifying the dihedral terms describing the $\chi$ angle. The current and relatively simple scheme for implementing stafix also reflected the need not to introduce too many off-diagonal terms in the LJ matrix as these could slow down the simulations on some machines and complicate the implementation. In fact, given the large number of overlaps with the protein atomic types that have to be resolved when applying stafix, we suggest the current version of stafix strikes a good balance between sophistication and practicality. As stafix is intended to be a goal-specific modification of the core simulation *ff*, the results are quite insensitive to details of the stafix implementation; i.e., any reasonable variant will provide the desired results.

**Formation of RNA globules and degradation of the interface in simulations of the bound HuR RRM3–5′-UUUUU-3′ complex.**

Experiments have shown that the RNA stably binds in pockets p2 and p3 of HuR RRM3 while binding in pockets p1 and p4 is disordered. (3) In simulations, this results into a characteristic behaviour where binding in pockets p2 and p3 is stable while numerous disruptions and reformations of binding within the other pockets are observed on the microsecond timescale. Although such behaviour is consistent with the experiments to an extent, (3) it makes the system extremely sensitive to accumulating spurious RNA self-interactions as these are clearly preferred by the standard OL3 *ff* over the native protein-RNA interactions. In other words, the dynamics of the p1 and p4 pocket which is correct per se and desirable exposes the system to formation of spurious self-interactions. Similarly to the free RNA, the problem is best described as drastic lowering of RNA's SAS as the molecule folds upon itself (main text Figure 4). Coupled with this is the size reduction of the protein-RNA interface as it is replaced by the competing RNA self-interactions. In the end, this leads to very distorted complex interface where binding in pockets p1 and p4 can no longer be reformed and only binding in pockets p2 and p3 is still maintained (main text Figure 4). The permanent formation of RNA self-interactions is entirely prevented when using OL3-stafix 0.5 or 0.1 (Figure S3). These simulations still reflect the experimentally documented disordered nature of RNA binding in pockets p1 and p4 as numerous and extensive disruptions are observed on the simulation timescale. However, elimination of the RNA self-interactions routinely allows their reformation which is far less likely with the standard OL3 *ff* (Figure S3).

**Substrate specificity of HuR RRM3 is not abolished with OL3-stafix.** We showed that it is possible to routinely observe spontaneous binding of the target RNA sequences to HuR RRM3 with OL3-stafix *ff* (main text). Although we consider this a very encouraging result, the stafix obviously represents a major modification of the balance between RNA-RNA, protein-RNA, and RNA-water interactions in favour of the latter two. As such, we needed to make sure that application of stafix does not allow non-target RNA sequences to bind with the protein. To verify this, we performed SBSs of a completely off-target 5′-CCCCC-3′ RNA. The poly-C RNA would transiently sample the pockets where target RNAs bind (Figure S5, #37-#40). This was expected as these pockets are lined with phenylalanine and tyrosine side-chains which can non-specifically interact with any nucleotide. (3,4) However, due to lack of compatible donor-acceptor groups which could form H-bonds, the binding was unstable and the RNA eventually disengaged from the native binding site. Afterwards, it continued to transiently sample various non-native sites on the protein's surface. In conclusion, we suggest that applying stafix does not abolish the substrate specificity of HuR RRM3.

**Excessively stable interaction between the sulphur of cysteine 245 and the RNA.**
In two of the unsuccessful SBS of the HuR RRM3 complex (Figure S5, #5 #18), we observed the RNA forming long-term interaction between one of the bases and the sulphur atom of cystein 245 which is part of the pocket p4. This interaction stalled the RNA in its conformational search and stopped it from assuming the native protein-RNA binding mode. Based on our previous study, (5) we suspect this to be an artefact of the AMBER LJ potential parameter of the sulphur atom which might not be ideal in all chemical contexts, producing excessively attractive nonbonded interactions. We subsequently verified that the RNA which already formed such interaction with sulphur in simulations can be 'freed' by changing the AMBER sulphur LJ parameter to the thiolate sulphur parameter from CHARMM. (6) The CHARMM sulphur parameter could be a superior choice in simulations, although its ultimate performance will also depend on the utilized water model and other factors.

**Comparison of the MD simulation ensembles of TNs and HN with experimental data – methodological part.**

We analyzed four NMR observables, i.e., (i) backbone $^3$J scalar couplings, (ii) sugar $^3$J scalar couplings, (iii) nuclear Overhauser effect intensities (NOEs), and (iv) the absence of specific peaks in NOE spectroscopy (unobserved NOEs – uNOEs). Their combination provided the total $\chi^2$ value. The lower $\chi^2$ value, the better agreement between the simulation ensemble and the experimental data ($\chi^2$ values below 1 indicate good agreement with the experiment). $^3$J scalar couplings were obtained from the Karplus relationships, while NOEs and uNOEs were obtained from MD simulations as averages over the N samples, i.e., $(u)NOE_{CALC} = \left(\frac{\sum_i^N r_i^{-6}}{N}\right)^{-1/6}$. Calculated NOEs were directly compared against experimental values. uNOE signals were considered as violated when the calculated value from the simulation was shorter than the assigned one from the experiment (see Ref. (7) for more details).

**Simulations of a fully bound SRSF1 RRM2–5′-AGGAC-3′ complex.**

Simulations started from a fully bound SRSF1 RRM2–5′-AGGAC-3′ complex yielded also significantly more stable protein-RNA interface with OL3-stafix *ff*. Namely, in two simulations using the standard OL3 *ff*, we observed the $A_4$ permanently detaching soon after start and after ~1.2 μs, respectively, while forming extensive self-interactions with neighboring nucleotides. In one of the simulations, the second pocket was also lost after ~5 μs. No such progressive degradation of the protein-RNA interface and RNA globule formation was seen in OL3-stafix simulations (Figure S8).

**SBS results when using the standard OL3 *ff*.** Out of the twelve SBSs of the HuR RRM3–5′-UUUUU-3′ system, we observed some binding in three simulations when stafix was not used (Figure S5, #26 #27 #31). In the first of those three, the binding occurred at the very beginning of the simulation before the RNA could extensively self-interact and form the globule. However, unlike in the stafix simulations, the binding was not stable on the entire simulation timescale as the RNA became gradually affected by the self-interactions, similarly as when starting with an already bound RNA. After 3.6 μs the native binding interface was largely lost in favour of the RNA self-interactions (Figure S5, #26). In the second partially successful SBS, we observed the RNA binding only at the very end (9.9 μs) of the simulation (Figure S5, #27). This SBS represent the sole case where the protein-RNA interactions managed to outcompete the RNA self-interactions but only after the RNA spent nearly 5 μs in the vicinity of the native binding pockets attempting to do so. Upon extension of this simulation, we observed that the native binding mode was once again disrupted by RNA self-interactions, similarly to when starting the simulation from a fully-bound structure. In the third partially successful SBS (Figure S5, #31), the RNA spontaneously bound after 1.8 μs but only pockets p2 and p3 were regularly occupied and there were frequent disruptions caused by the RNA self-interactions. Therefore, although a few simulations without stafix have shown some tendency for binding, no binding event was really complete and successful.

The difference was even larger for the HuR RRM3–5′-UUUA-3′ system where none of the SBSs using the standard OL3 *ff* showed any sign of binding (Figure S5, #41–#46) while the RNA successfully bound in all OL3-stafix SBSs (Figure S5, #47–#51). The SBSs using the standard OL3 *ff* were also completely unsuccessful for the SRSF1 RRM2–5′-AGGAC-3′ system (Figure S6, #13-#18 ) as a result of extensive formation of RNA globules (Figure S7).

**OL3-stafix of TNs and HN.**

In addition to the protein-RNA complexes, we also tested performance of the stafix on commonly studied isolated ssRNA motifs – tetranucleotides and hexanucleotide (Table S1).

For the TNs, the stafix abolishes both the spurious intercalated and native A-form-like conformations in simulations, especially at higher scaling factors (Table S2). In other words, its main effect is allowing the TNs to dominantly sample the unstructured single-stranded conformation; an effect which is of significant benefit in protein-ssRNA simulations as it both stabilizes the protein-RNA interface and allows observing spontaneous complex formation (see the main text). The resulting ensemble is not in full agreement with the NMR data (Table S2), although we reiterate that this was never the goal of stafix. A notable exception was the r(UUUU) TN which showed improvement when using stafix 0.5, producing ensemble with $^3$J-couplings very close to experimental values (Table S2). This indicates that weakening of the vdW interactions and disordering the ensemble by stafix may bring some advantage for the r(UUUU) description; note that it is difficult to characterize the r(UUUU) ensemble based on the primary experimental data. Except of the r(UUUU), all the TNs and the HN showed closest agreement with the NMR data in stafix 0.75 simulations with simultaneous use of gHBfix and tHBfix potentials. (2,7) The performance was also mildly improved compared to the standard OL3 *ff* as use of stafix 0.75 helped eliminating the intercalated states. At even milder scaling factors, stafix could be of potential interest also for simulations of free short ssRNAs and possibly other motifs, as it helps eliminating excessive stacking, especially when combined with gHBfix and tHBfix potentials which adjust stability of H-bonds. (2,7)

**Benchmark simulations of the HuR RRM3 using three different protein *ff*s rationalize the advantage of ff12SB.**

In all our simulations presented in the main text (main text Table 1), we used the ff12SB *ff* to describe the proteins. The reason for preferring this protein *ff* version instead of the newer ff14SB or ff19SB is the superior description of the phenylalanine and tyrosine side-chains provided by ff12SB. Namely, both ff14SB and ff19SB are missing one specific dihedral term which is present in ff12SB and which introduces an increased energy barrier for $\chi_2$ side-chain rotation. As a result, in our simulations of the isolated HuR RRM3 protein, we observed similar distribution of the $\chi_2$ dihedrals of the phenylalanine residues with all three protein *ffs* but significantly slower rotational transitions with ff12SB. For example, for the surface exposed F279 residue, this corresponded to an average time between rotations of ca 22 ns with ff12SB but only 1 ns with ff14SB and ff19SB (Figure S12). The fast rotation of the aromatic side-chains can cause problems in simulations of protein-RNA complexes as these amino acids often form part of the protein-RNA interfaces (8) and are especially prominent in RRM domains studied in this work. (3,9) As a result, the ff12SB produces ensembles of the HuR RRM3 which are more in agreement with the experimental data (3,9) (Figure S12) and we suggest it is the more appropriate choice for simulations of the RRM protein-RNA complexes in general. A tempting option would be to straightforwardly excise the dihedral term responsible for the better performance of ff12SB and apply it as a modification of the ff14SB and ff19SB *ff*s. However, extensive testing would be required to identify possible side-effects of such approach.

**OL3-stafix simulations of A-RNA duplex.**

In simulations of the A-RNA duplex, we tested stafix factors 0.5 and 0.75 and compared their simulation performance against the standard OL3 *ff*. The effects of OL3-stafix 0.75 on A-RNA helical parameters were almost exactly midway between standard OL3 *ff* and OL3-stafix 0.5. The most significant change in A-RNA geometry caused by stafix was the decrease of propeller twist, which can be related to the reduced base stacking, since propeller twist increases overlap of bases in the intra-strand stacking. We also observed decreased inclination, increased slide and 2-4° reduction of the $\chi$ dihedral (Table S3). There were also increased fluctuations of AU base pairs and occurrence of short-lived water-mediated states

(Figure S10). We also observed faster fraying of the terminal base pairs, which, however can also be seen with standard OL3 *ff* (Figure S11). In conclusion, changes to the A-RNA duplex geometry in OL3-stafix simulations are surprisingly minor compared to the standard OL3 *ff*, especially when one considers how extensively stafix alters the system's potential energy function. There were no signs of the helical structure collapsing even with stafix scaling of 0.5. Therefore, stafix could potentially be also applied for protein-RNA systems where RNA stem-loops are bound and the loop part of the RNA forms the protein-RNA interface (e.g. the U1A RRM protein-RNA complex). We, however, again reiterate that the stafix modification is not intended for simulations of folded RNAs. However, in protein-RNA binding studies, its applicability could be broader than just simulations with short ssRNA segments.

#### Supplementary Tables

Table S1. List of all MD simulations of the tetranucleotides (TN), hexanucleotide (HN), A-RNA duplex, isolated HuR RRM3 protein, and free r(UUUUU).[a]

| RNA Motif | *ff* modification [b] | water model | simulations × length [μs] |
|---|---|---|---|
| **Target sequence of HuR RRM3** | | | |
| r(UUUUU) | - | SPC/E | 1 × 2 |
| r(UUUUU) | stafix 0.50 | SPC/E | 2 × 2[c] |
| r(UUUUU) | stafix 0.10 | SPC/E | 1 × 2 |
| r(UUUUU) | - | OPC | 1 × 2 |
| r(UUUUU) | Garcia *ff* [d] | SPC/E | 1 × 2 |
| r(UUUUU) | DESRES *ff* [d] | TIP4PD | 1 × 2 |
| **TNs and HN[b]** | | | |
| r(AAAA) | stafix 0.25 | SPC/E | 1 × 10 |
| r(AAAA) | stafix 0.50 | SPC/E | 1 × 10 |
| r(AAAA) | stafix 0.75 | SPC/E | 1 × 10 |
| r(AAAA) | stafix 0.75, gHBfix, tHBfix | SPC/E | 1 × 10 |
| r(CAAU) | stafix 0.25 | SPC/E | 1 × 10 |
| r(CAAU) | stafix 0.50 | SPC/E | 1 × 10 |
| r(CAAU) | stafix 0.75 | SPC/E | 1 × 10 |
| r(CAAU) | stafix 0.75, gHBfix, tHBfix | SPC/E | 1 × 10 |
| r(CAAU) | stafix 0.25 | OPC | 1 × 10 |
| r(CAAU) | stafix 0.50 | OPC | 1 × 10 |
| r(CAAU) | stafix 0.75 | OPC | 1 × 10 |
| r(CAAU) | stafix 0.75, gHBfix, tHBfix | OPC | 1 × 10 |
| r(CCCC) | stafix 0.25 | SPC/E | 1 × 1 |
| r(CCCC) | stafix 0.50 | SPC/E | 1 × 1 |
| r(CCCC) | stafix 0.75 | SPC/E | 1 × 1 |
| r(CCCC) | - | SPC/E | 1 × 1 |
| r(UUUU) | stafix 0.25 | SPC/E | 1 × 10 |
| r(UUUU) | stafix 0.50 | SPC/E | 1 × 10 |
| r(UUUU) | stafix 0.75 | SPC/E | 1 × 10 |
| r(UUUU) | stafix 0.75, gHBfix, tHBfix | SPC/E | 1 × 10 |

| | | | |
|---|---|---|---|
| r(UCAAUC) | stafix 0.25 | SPC/E | 1 × 10 |
| r(UCAAUC) | stafix 0.50 | SPC/E | 1 × 10 |
| r(UCAAUC) | stafix 0.75 | SPC/E | 1 × 10 |
| r(UCAAUC) | stafix 0.75, gHBfix, tHBfix | SPC/E | 1 × 10 |
| **A-RNA duplex**[e] | | | |
| duplex | HBfix (terminal bps) | SPC/E | 1 × 1 |
| duplex | stafix 0.75, HBfix (terminal bps) | SPC/E | 1 × 1 |
| duplex | stafix 0.50, HBfix (terminal bps) | SPC/E | 1 × 3 |
| duplex | stafix 0.50 | SPC/E | 1 × 1 |
| duplex | - | SPC/E | 2 × 1 |
| **Isolated HuR RRM3** | | | |
| **protein *ff*** | **Water model** | **simulations × length [μs]** | |
| ff12SB | SPC/E | 1 × 10 | |
| ff14SB | SPC/E | 1 × 10 | |
| ff19SB | OPC | 1 × 10 | |

[a]Simulations of r(UUUUU) in OPC water, the TNs and the HN were run with the $\chi$OL3$_{CP}$ RNA force field (10,11) instead of the standard OL3. For all other details, see the main text Methods.

[b]Some of the stafix 0.75 simulations were also tested together with the gHBfix (version known also as gHBfix19) (2) and tHBfix (7) potentials. tHBfix settings for each motif were taken from our previous work. (7) When nothing is noted, the basic (core) OL3 *ff* is used.

[c]One of the simulations was started from the end of the standard OL3 *ff* simulation in which an RNA globule already formed.

[d]Simulations utilizing the alternative RNA *ff*s proposed by Chen&Garcia (12) (García *ff*) and Tan et al. (DESRES *ff*) (13), respectively. For the DESRES simulation, the suggested CHARMM parameters for ions were utilized along with the recommended TIP4PD water model.

[e]Sequence of the A-RNA duplex was r(CGCGGGAAAUCCCGCG). In some of the simulations, a 2 kcal/mol structure-specific HBfix (1) was used to stabilize the Watson-Crick H-Bonds of the terminal base pairs.

Table S2. Conformational analyses of RNA TNs and HN in MD simulations and their comparison with experiments.[a]

| **RNA Motif** | ***ff* modification[a]** | **$\chi^2$ [$^3$J-backbone, $^3$J-sugar, NOE, uNOE / Total] [b]** | **Clustering [%] [A-form-like / Intercalated / 1-3_2-4 Stack / Unstructured single strand]** |
|---|---|---|---|
| r(AAAA) | stafix 0.25 | 1.29, 9.79, 12.46, 0.33 / **2.14** | ~0/~0/~0/~99 |
| r(AAAA) | stafix 0.50 | 1.10, 3.92, 4.81, 0.44 / **1.11** | ~0/~0/~0/~97 |
| r(AAAA) | stafix 0.75 | 1.07, 1.79, 2.50, 1.76 / **1.81** | ~77/~8/~0/~0 |
| r(AAAA) | stafix 0.75, gHBfix, tHBfix | 0.89, 2.96, 2.97, 0.80 / **1.14** | ~76/~0/~3/~0 |
| r(CAAU) | stafix 0.25 | 1.31, 9.10, 17.30, 3.09 / **4.42** | ~0/~0/~0/~99 |
| r(CAAU) | stafix 0.50 | 1.15, 3.44, 7.37, 3.33 / **3.61** | ~0/~0/~0/~98 |
| r(CAAU) | stafix 0.75 | 1.15, 3.44, 4.50, 6.37 / **5.93** | ~78/~18/~0/ ~0 |
| r(CAAU) | stafix 0.75, gHBfix, tHBfix | 0.89, 2.35, 4.14, 3.69 / **3.59** | ~95/~0/~0/~0 |
| r(CAAU)[c] | stafix 0.25 | 1.33, 10.26, 20.95, 3.19 / **4.86** | ~0/~0/~0/~100 |
| r(CAAU)[c] | stafix 0.50 | 1.13, 4.61, 8.46, 2.76 / **3.24** | ~0/~0/~0/~98 |
| r(CAAU)[c] | stafix 0.75 | 1.02, 1.79, 3.85, 5.06 / **4.72** | ~90/~7/~0/~0 |
| r(CAAU)[c] | stafix 0.75, gHBfix, tHBfix | 0.81, 2.04, 3.52, 2.71 / **2.69** | ~95/~0/~0/~0 |
| r(CCCC) | stafix 0.25 | 1.85, 7.94, 22.31, 3.27 / **5.09** | ~0/~0/~0/~98 |
| r(CCCC) | stafix 0.50 | 1.42, 4.50, 8.39, 2.65 / **3.18** | ~0/~0/~0/~96 |
| r(CCCC) | stafix 0.75 | 1.06, 1.98, 5.43, 4.12 / **4.00** | ~33/~62/~0/~1 |
| r(CCCC) | - | 1.13, 0.66, 7.33, 5.29 / **5.10** | ~18/~79/~0/~0 |
| r(UUUU) | stafix 0.25 | 0.97, 2.49, 3.63, 0.64 / **0.80** | ~0/~0/~0/~99 |
| r(UUUU) | stafix 0.50 | 0.84, 0.96, 3.85, 0.42 / **0.56** | ~0/~0/~0/~98 |
| r(UUUU) | stafix 0.75 | 0.76, 1.26, 4.55, 0.65 / **0.79** | ~96/~0/~0/~0 |
| r(UUUU) | stafix 0.75, gHBfix, tHBfix | 0.65, 1.05, 3.84, 0.70 / **0.80** | ~0/~0/~4/~91 |
| r(UCAAUC) | stafix 0.25 | 1.02, 10.92, 13.93, 1.11 / **2.72** | ~0/~0/~0/~85 |
| r(UCAAUC) | stafix 0.50 | 0.97, 4.82, 7.80, 0.72 / **1.59** | ~0/~0/~0/~82 |
| r(UCAAUC) | stafix 0.75 | 0.96, 2.79, 6.17, 0.62 | ~79/~2/~0/~2 |

| | | | |
|---|---|---|---|
| | | / **1.29** | |
| r(UCAAUC) | stafix 0.75, gHBfix, tHBfix | 0.70, 3.29, 5.36, 0.65 / **1.23** | ~79/~0/~0/~0 |

[a]See footnote b of Table S1.
[b] $\chi^2$ values were obtained by comparing calculated and experimental backbone $^3J$ scalar couplings, sugar $^3J$ scalar couplings, nuclear Overhauser effect intensities (NOEs), and the absence of specific NOE peaks (uNOEs, see above and Ref. (7) for details).
[c]Simulations were done with the OPC water model.

Table S3. Average values of selected helical parameters in MD simulations of the A-RNA duplex.

| **parameter** | **stafix factor** | **average (±)** [˚] |
|---|---|---|
| propeller (AU pairs) | no stafix | -11.9 (8.3) |
| | 0.75 | -10.1 (8.7) |
| | 0.50 | -7.6 (9.0) |
| propeller (GC pairs) | no stafix | -11.4 (7.5) |
| | 0.75 | -9.0 (8.0) |
| | 0.50 | -6.2 (8.6) |
| inclination | no stafix | 18 (11) |
| | 0.75 | 16 (12) |
| | 0.50 | 15 (12) |
| **parameter** | **stafix factor** | **average (±)** [Å] |
| slide | no stafix | -1.6 (0.5) |
| | 0.75 | -1.7 (0.5) |
| | 0.50 | -1.8 (0.5) |

**Supplementary Figures**

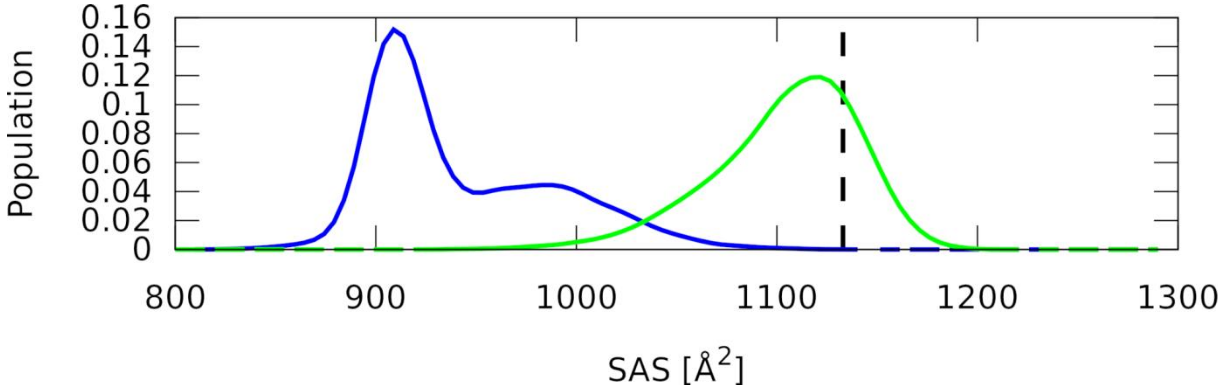

Figure S1. **MD simulations of the free 5′-UUUA-3′ RNA.** Comparison of the RNA SAS values observed with standard OL3 *ff* (blue) and in OL3-stafix 0.1 simulations (green). The vertical dashed line indicates SAS value of the HuR RRM3-bound conformation.

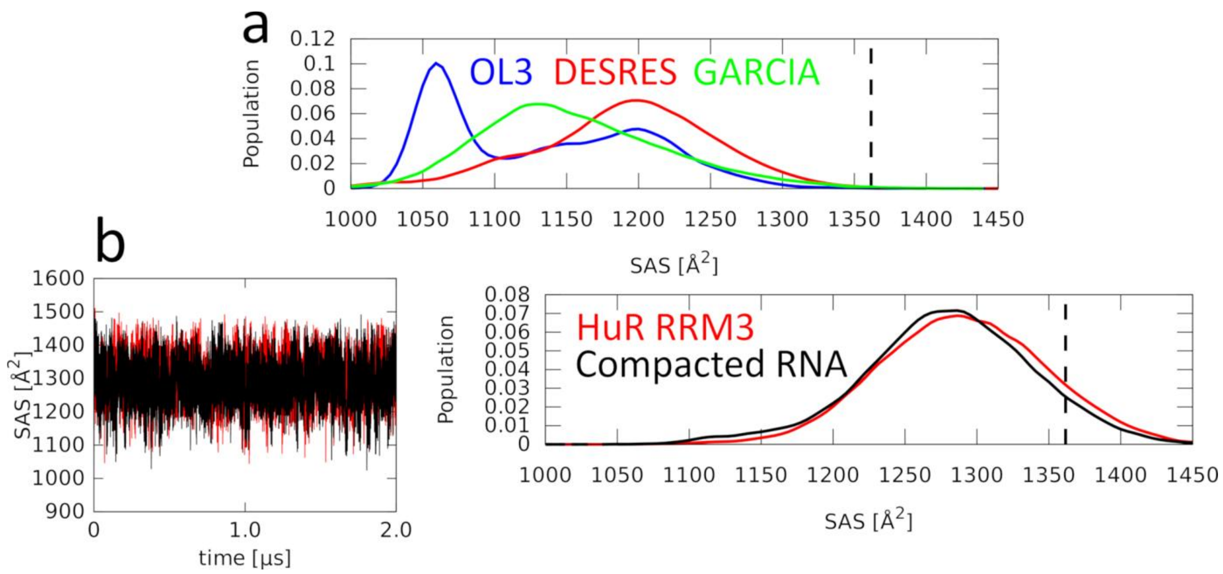

Figure S2. **MD simulations of the free 5′-UUUUU-3′ RNA with alternative RNA *ff*s or starting structures.** a) Histograms of the RNA solvent-accessible surface (SAS) when simulating with different RNA *ff*s – OL3, (14) DESRES, (13) and García. (12) All three tested RNA force fields fail to produce 5′-UUUUU-3′ RNA ensembles with SAS value of the HuR RRM3-bound conformation (the vertical dashed line) accessible. b) c) Comparison of the SAS values observed in OL3-stafix 0.5 simulations when starting from HuR RRM3-bound conformation (red) or from an RNA globule structure produced by the standard OL3 *ff*. The use of stafix melts the globule within first few nanoseconds. The resulting ensembles are then essentially identical within the limits of the sampling.

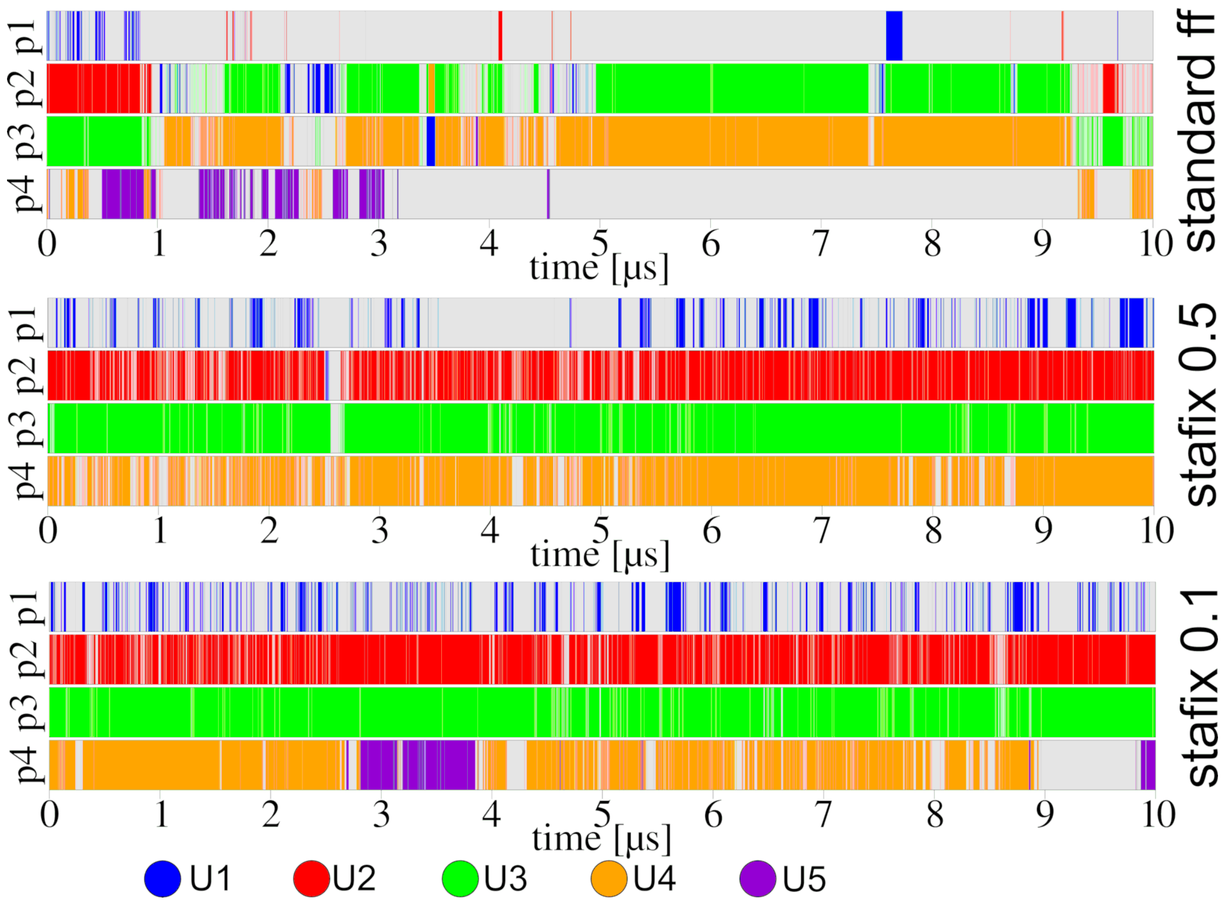

Figure S3. **Time-development of the binding pocket (p1-p4) occupancies by specific nucleotides in selected simulations of the fully bound HuR RRM3–5′-UUUUU-3′ complex.** Without stafix, we observe gradual and ultimately irreversible degradation of the complex interface as the RNA collapses into a globule. The stafix 0.1 trajectory nicely captures competition of U4 and U5 for the p4 pocket. Note that the binding-occupancy descriptor (see the main text Methods for definition) does not visualise the full richness of the dynamics.

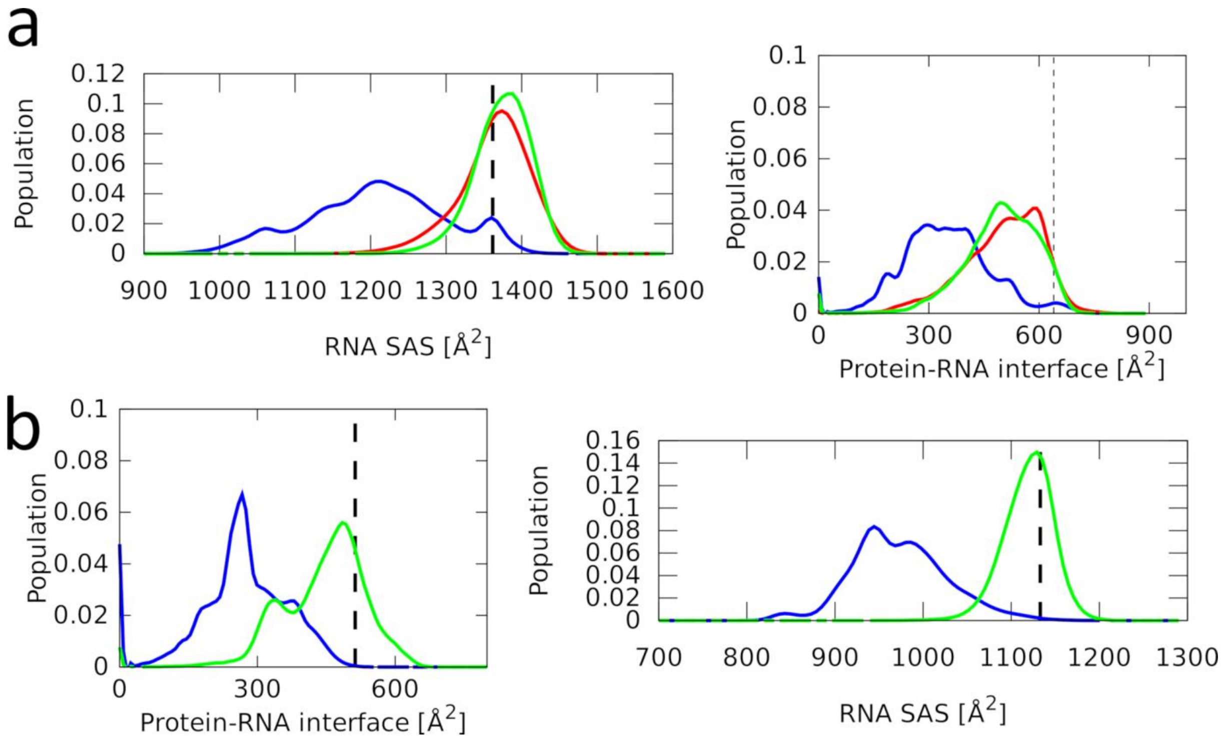

Figure S4. **SBSs of the HuR RRM3 5′-UUUUU-3′ (a) and 5′-UUUA-3′ (b) complexes.** a) b) Histograms of the RNA SAS and protein-RNA interface size, showing the RNA collapsing into a globule and limiting its contact with the protein in favour of RNA self-interactions with the standard OL3 *ff*. This can be entirely prevented with stafix. The vertical black dashed lines indicate the values in the native protein-RNA complex. The standard OL3 *ff* simulation is shown with blue curve while OL3 stafix 0.5 and 0.1 simulations are in red and green, respectively. The analysis was performed on combined simulation ensembles.

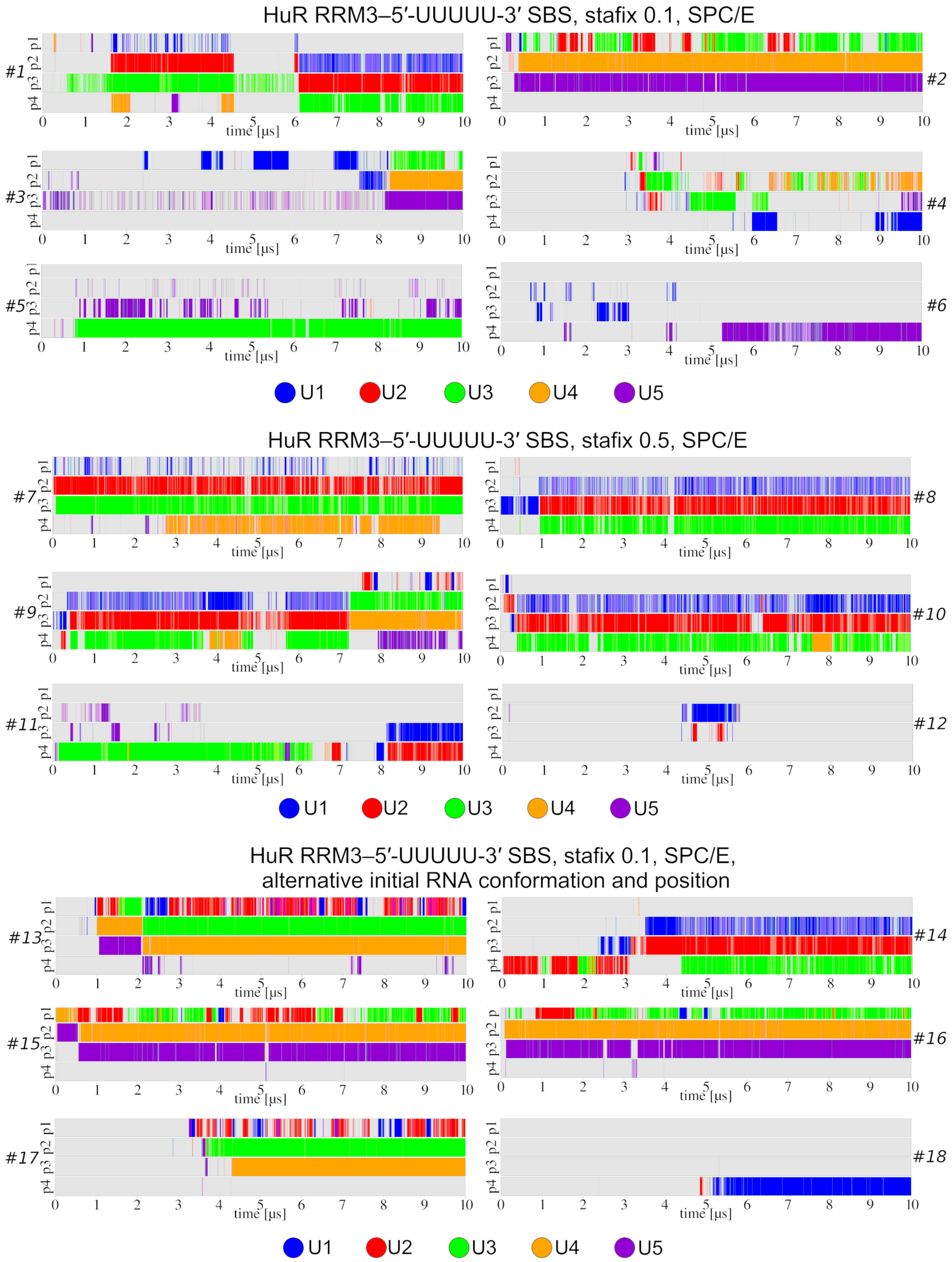

HuR RRM3–5′-UUUUU-3′ SBS, stafix 0.1, SPC/E
#1
#2
#3
#4
#5
#6
p1 p2 p3 p4
time [μs]
U1
U2
U3
U4
U5
HuR RRM3–5′-UUUUU-3′ SBS, stafix 0.5, SPC/E
#7
#8
#9
#10
#11
#12
p1 p2 p3 p4
time [μs]
U1
U2
U3
U4
U5
HuR RRM3–5′-UUUUU-3′ SBS, stafix 0.1, SPC/E,
alternative initial RNA conformation and position
#13
#14
#15
#16
#17
#18
p1 p2 p3 p4
time [μs]
U1
U2
U3
U4
U5

HuR RRM3–5′-UUUUU-3′ SBS, stafix 0.5, OPC

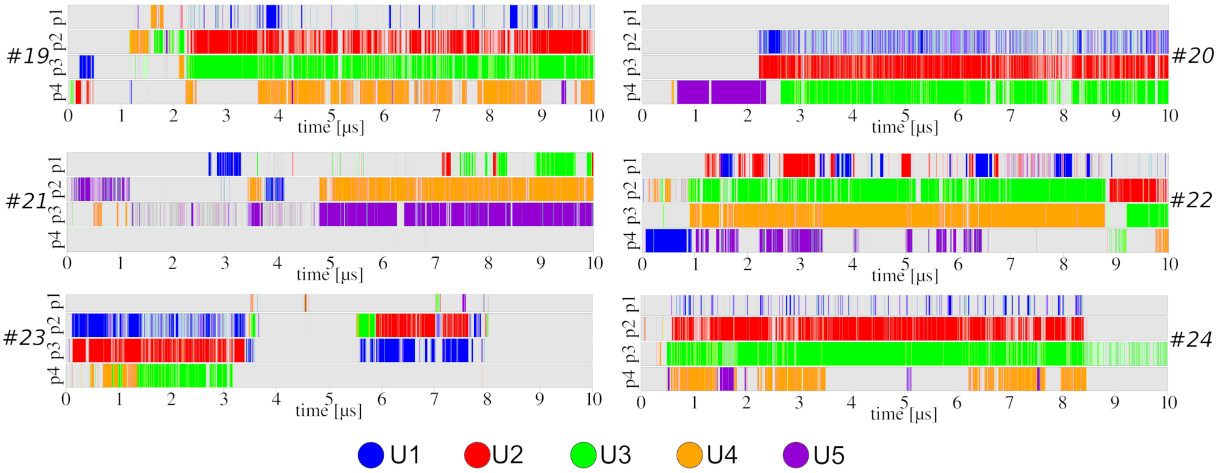

HuR RRM3–5′-UUUUU-3′ SBS, standard *ff*, SPC/E

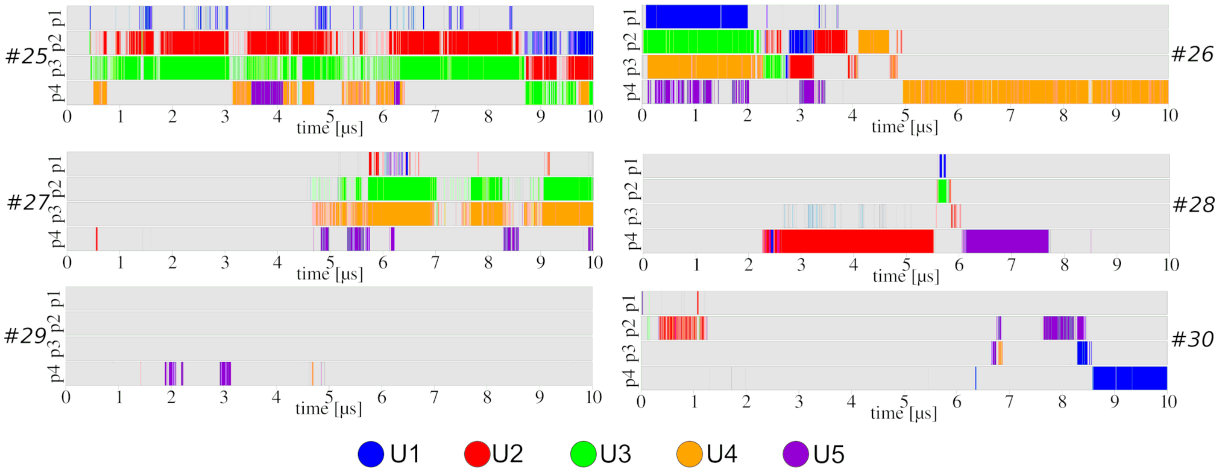

HuR RRM3–5′-UUUUU-3′ SBS, standard *ff*,OPC

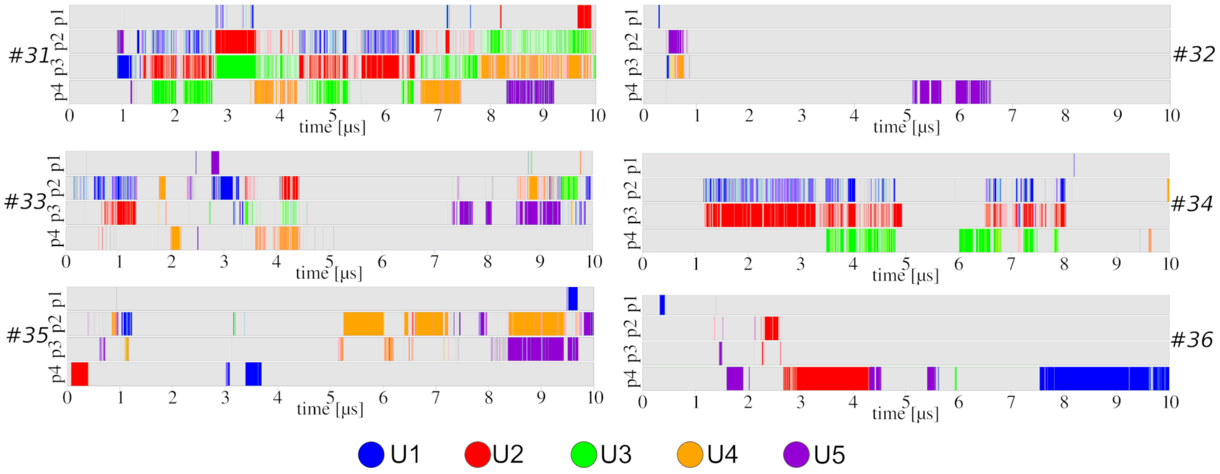

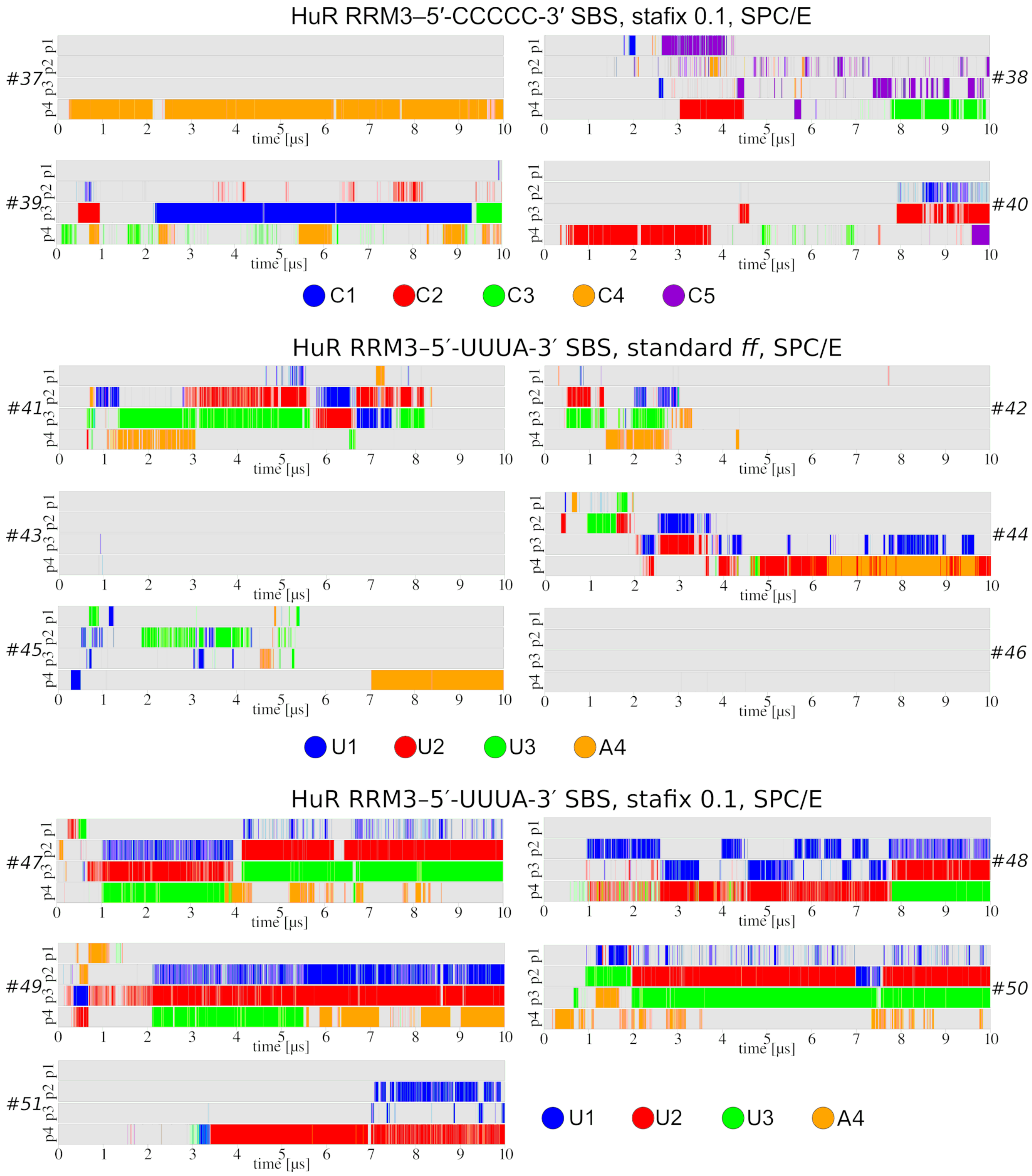

Figure S5. **Time-development of the binding pocket (p1-p4) occupancies by specific nucleotides in all SBS simulations of the HuR RRM3.** Each trajectory is assigned a unique identifier (#1 – #51).

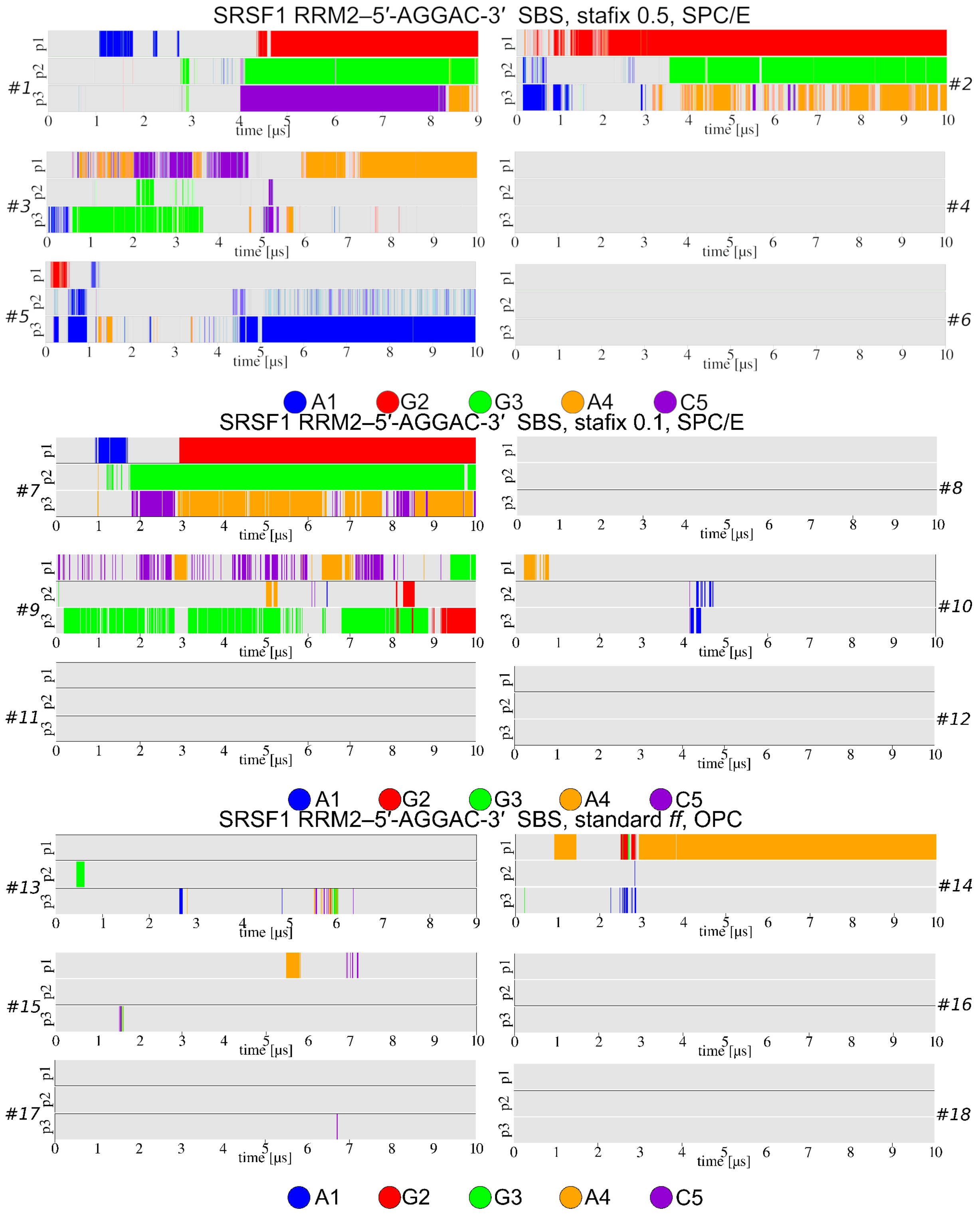

SRSF1 RRM2–5′-AGGAC-3′ SBS, stafix 0.5, SPC/E
#1
#2
#3
#4
#5
#6
p1
p2
p3
time [μs]
A1
G2
G3
A4
C5
SRSF1 RRM2–5′-AGGAC-3′ SBS, stafix 0.1, SPC/E
#7
#8
#9
#10
#11
#12
SRSF1 RRM2–5′-AGGAC-3′ SBS, standard ff, OPC
#13
#14
#15
#16
#17
#18

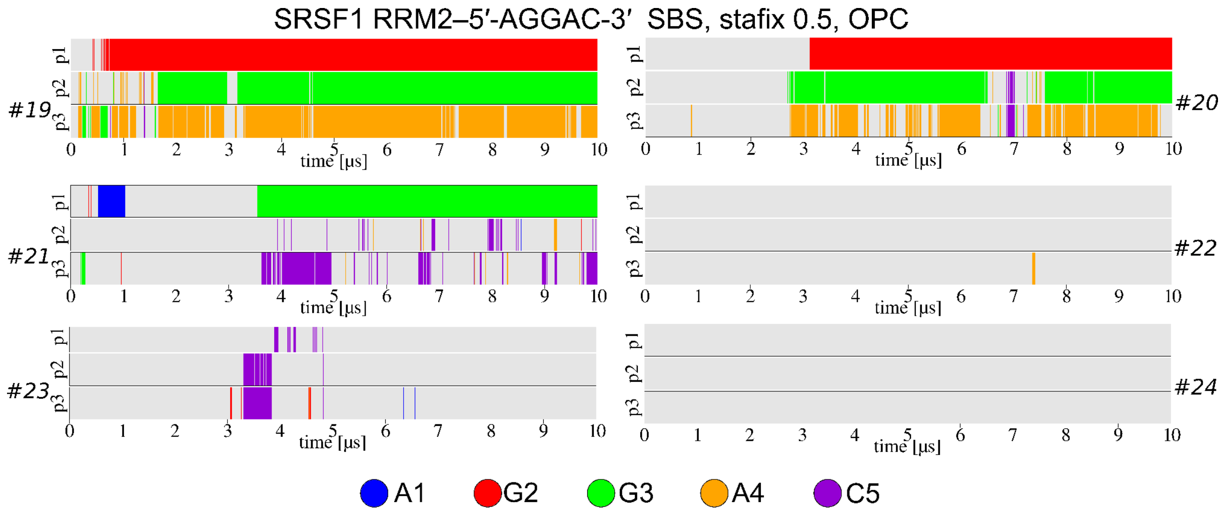

Figure S6. **Time-development of the binding pocket (p1-p3) occupancies by specific nucleotides in SBS simulations of the SRSF1 RRM2.** Each trajectory is assigned a unique identifier (#1 – #24).

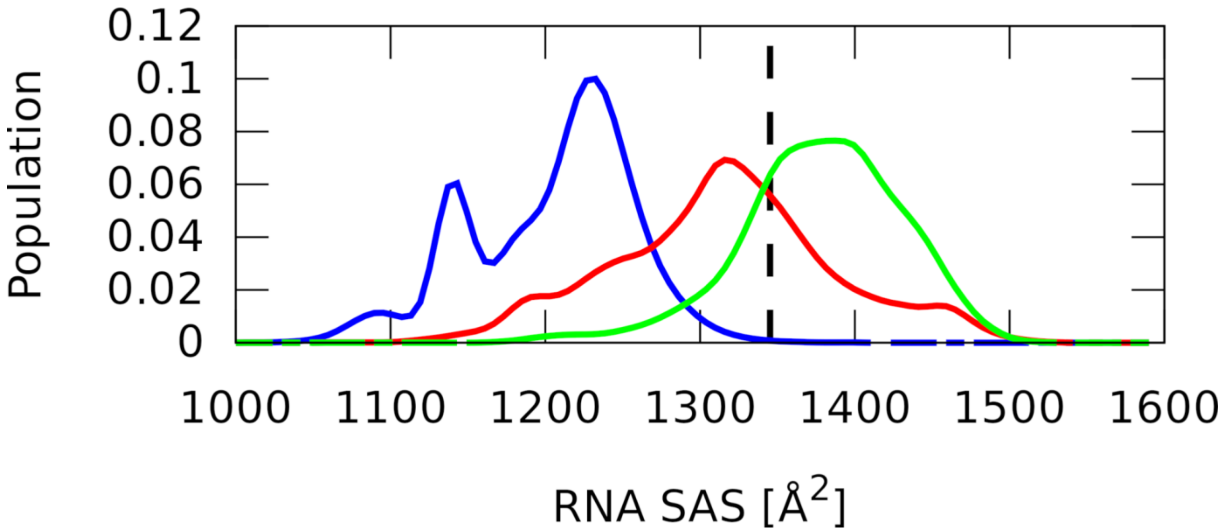

Figure S7. **Solvent-accessible surface of the 5′-AGGAC-3′ RNA in SBSs of the SRSF1 RRM2 complex.** The vertical black dashed line indicates the value corresponding to the native protein-RNA complex. The standard OL3 *ff* simulation is shown with blue curve while OL3 stafix 0.5 and 0.1 simulations are in red and green, respectively. Similarly to the target RNA in the HuR RRM3 complex, the SAS value seen in this complex is also not being sampled by the standard OL3 *ff*. The analysis was performed on combined simulation ensembles.

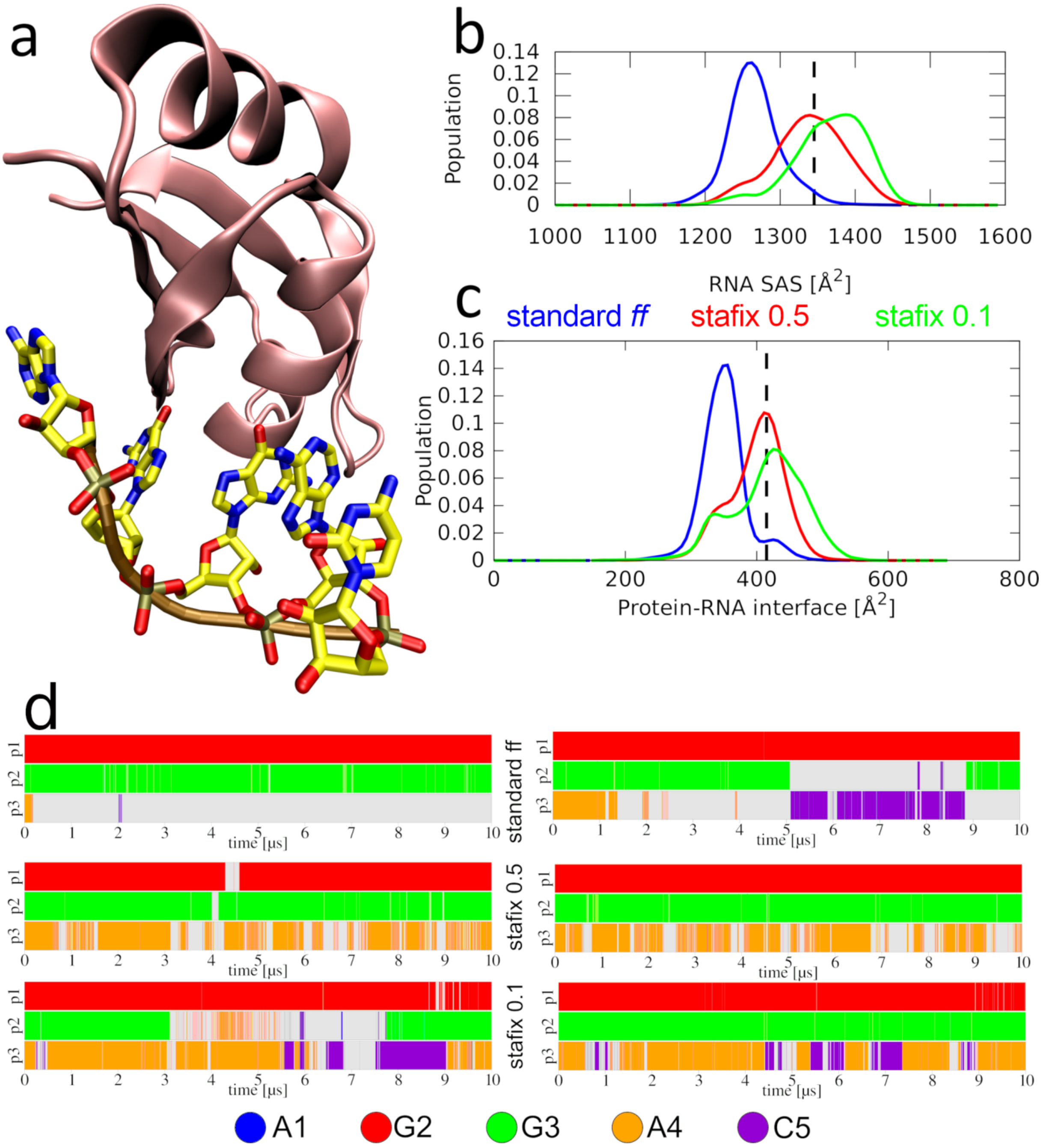

Figure S8. **MD simulations of the SRSF1 RRM2–5′-AGGAC-3′ complex starting from the bound state.** a) Snapshot from simulation using the standard OL3 *ff*. Without stafix, the $A_4$ in pocket p3 eventually detaches and forms a permanent triple base stack with the preceding guanosine and succeeding cytidine. b) Histograms of the RNA SAS. c) Histograms of the protein-RNA interface size. Panels b and c excellently show that the formation of RNA globules proceeds at the expense of the complex interface. The vertical black dashed lines indicate values corresponding to the native protein-RNA complex. d) Time-development of the binding pocket (p1-p3) occupancies by specific nucleotides.

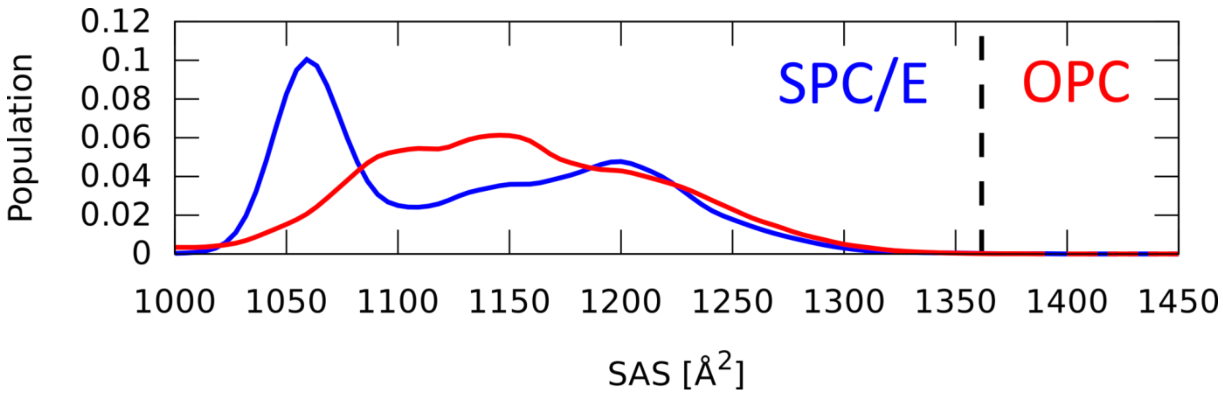

Figure S9. **Solvent-accessible surface of the free 5′-UUUUU-3′ RNA with different water models in standard OL3 *ff* simulations.** The vertical black dashed line indicates SAS value corresponding to the native protein-RNA complex. Both water models give similar SAS ranges and neither sample values seen in the complex, albeit the OPC model somewhat reduces the compactness.

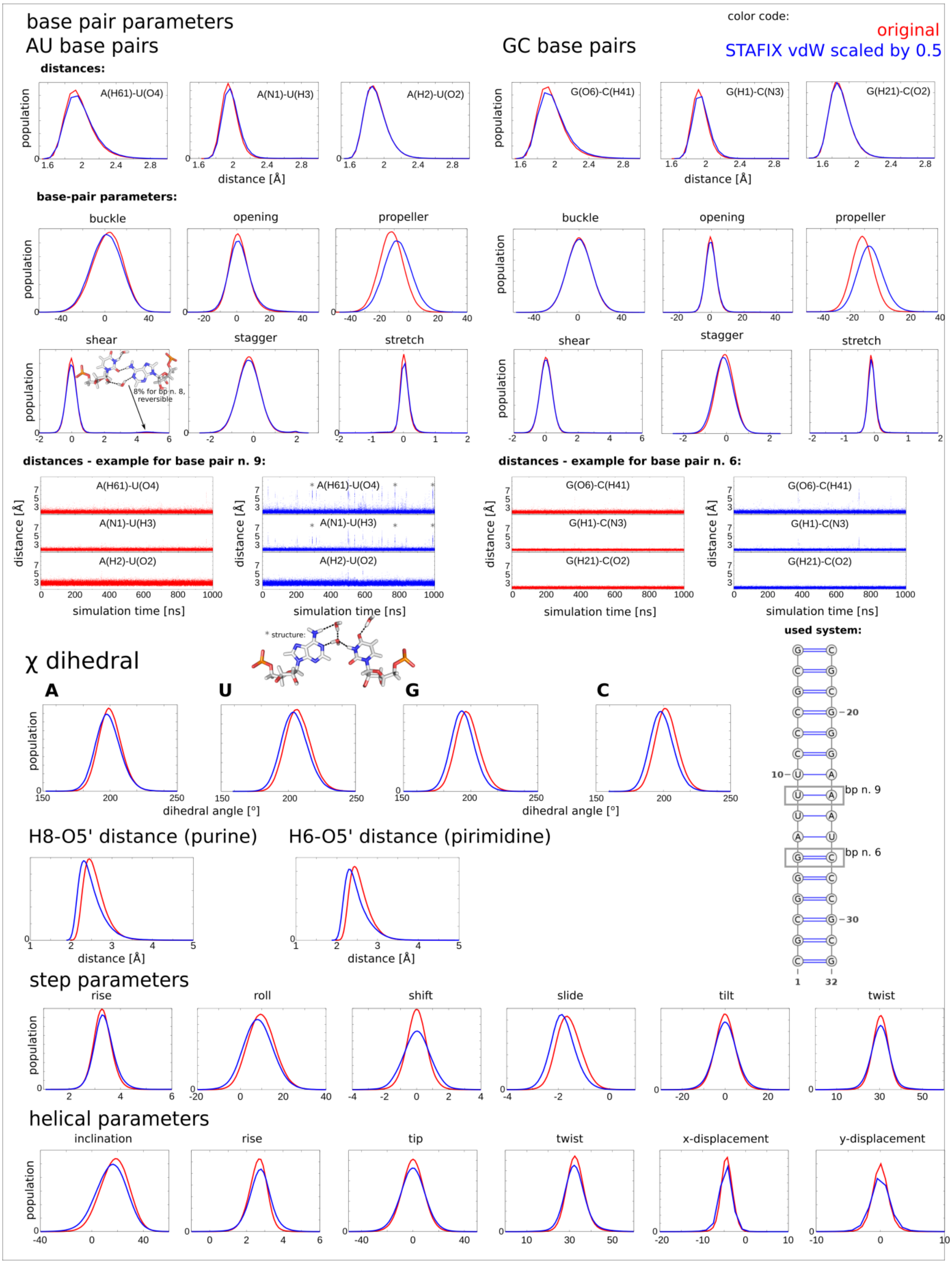

Figure S10. Histogram analyses of the base pair and helical parameters of A-RNA duplex as observed in selected MD simulations where terminal base pairs were stabilized with structure specific HBfix. Data for the standard *ff* and stafix with 0.5 rescaling factor is shown in red and blue, respectively. We do not show the stafix 0.75 data. The base-pair parameter values are based on aggregated data from all AU or GC pairs in the helix except the terminal base pairs. Analogously, base-pair step and helical parameter values are aggregated from all base-steps except the terminal ones. The 5% population of the sheared AU pair shown in the figure corresponds to reversible short-lived states (up to tens of nanoseconds).

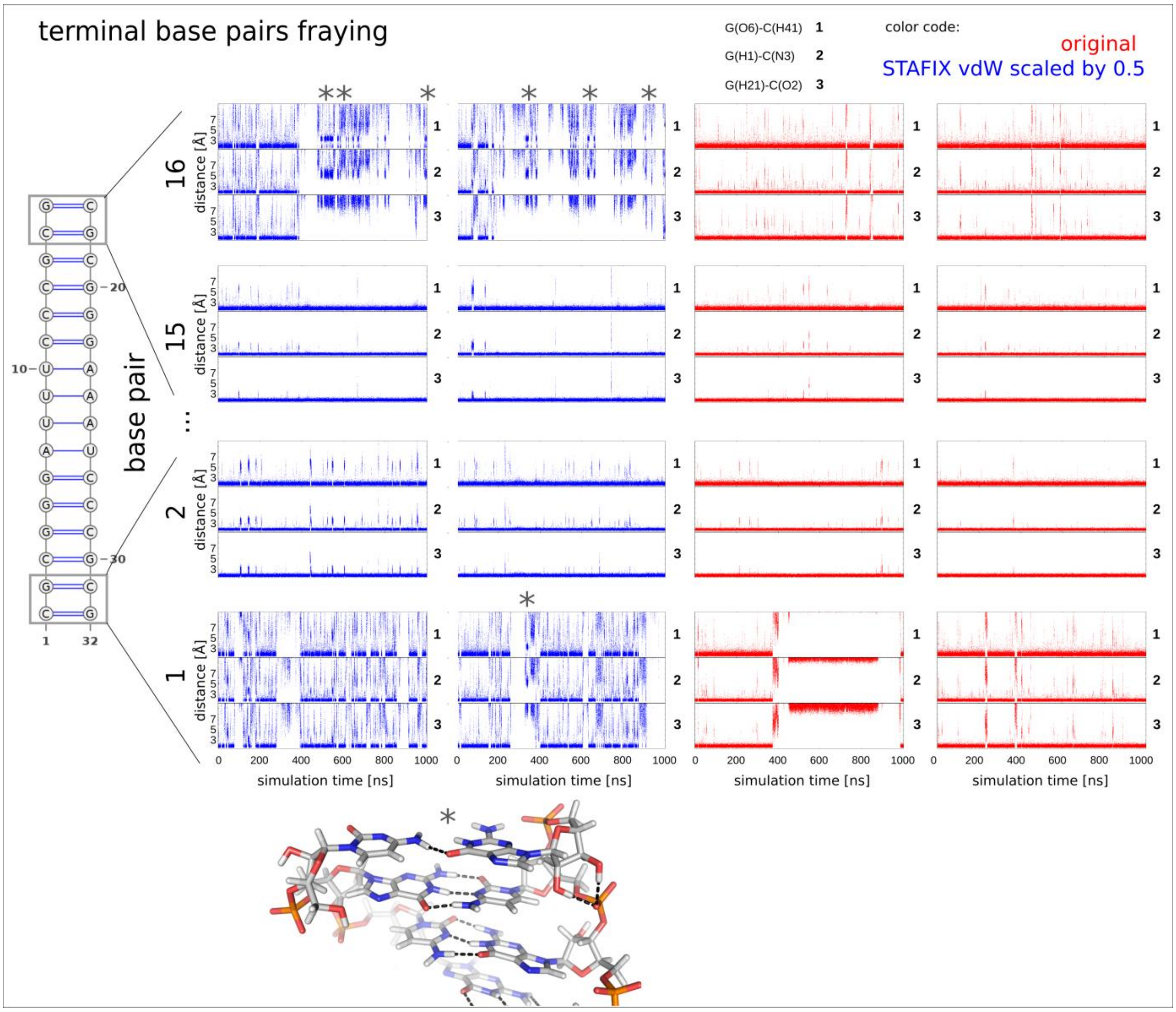

Figure S11. Time development of H-bonding distances of terminal and penultimate base pairs in A-RNA duplex simulations without HBfix. The "*" indicates specific terminal base pair conformation (bottom) sometimes seen in labeled trajectory parts.

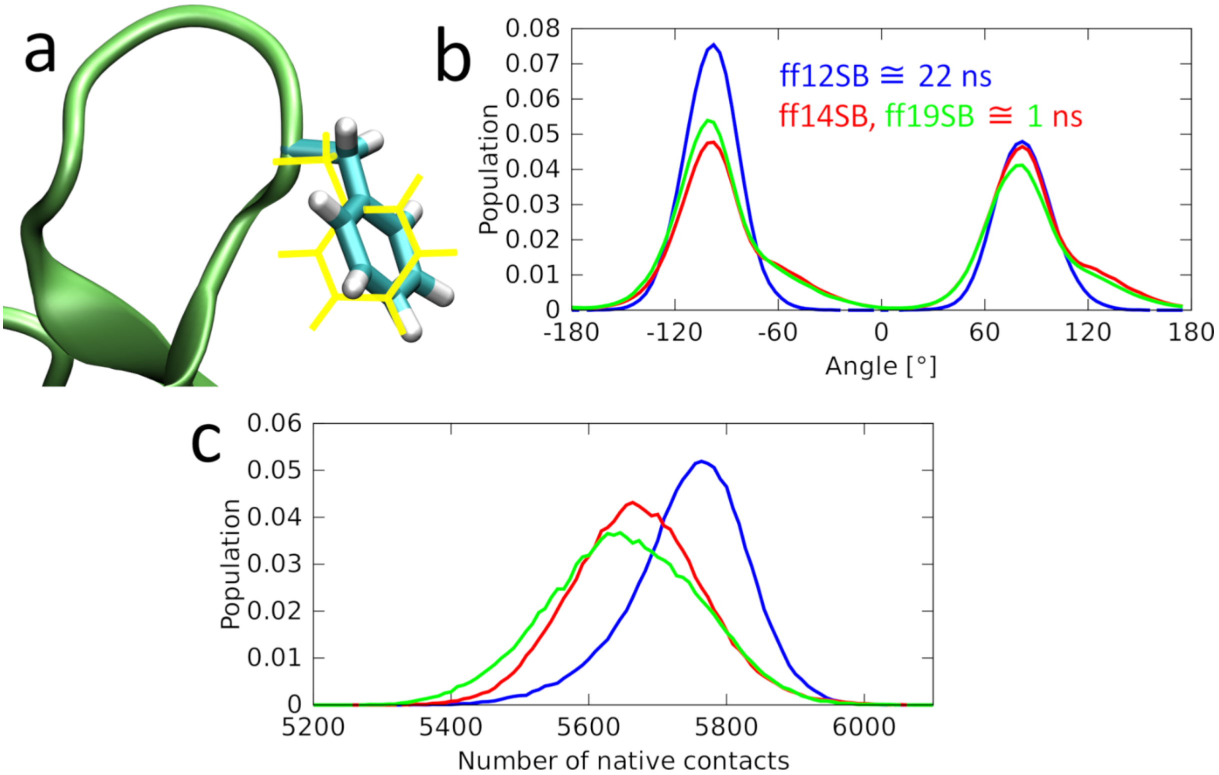

Figure S12. **Comparison between ff12SB, ff14SB, and ff19SB protein force fields.** a) Both rotamers of the phenylalanine side-chain are chemically equivalent. However, the transition state (shown in yellow) can cause disruptions of the surrounding segments. b) Histograms of the $\chi 2$ dihedral populations of F294 in simulations of the isolated HuR RRM3. Although all three force fields show similar distributions, the transitions (and therefore lifetimes of the individual rotamers) are more than twenty times faster with ff14SB and ff19SB than with the ff12SB. The frequent transitions can disrupt the structure of the protein-RNA complex. c) Histograms of the native contacts population. The graph indicates how many of the interatomic contacts present in the experimental structure (i.e. the native contacts) 'survive' during the MD simulation. The ff12SB force field maintains the native structure of the HuR RRM3 protein much better than the ff14SB or ff19SB force fields.
