## Supplementary material for "Spontaneous binding of single-stranded RNAs to RRM proteins visualised by unbiased atomistic simulations with rescaled RNA force field": Script

# this is script for scaling amber topology file using STAFIX scheme. parmed is used for scaling and must be located in your path.
# written by Pavlina Pokorna, 2022. For questions, contact Dr. Miroslav Krepl,
# Extensively tested with AMBER OL3 RNA ff as implemented in AMBER 14-22 and with ff12SB/ff14SB protein ffs. Also briefly tested with ff19SB protein ff. Works with STANDARD RNA residues.

import os
import sys
from math import sqrt
from distutils.spawn import find_executable


def is_to_be_scaled(resname,atname):
	scaling=False
	if resname in residue_namesA and atname not in atom_namesA+atom_namesSugar:
		scaling=True
	elif resname in residue_namesG and atname not in atom_namesG+atom_namesSugar:
		scaling=True
	elif resname in residue_namesC and atname not in atom_namesC+atom_namesSugar:
		scaling=True
	elif resname in residue_namesU and atname not in atom_namesU+atom_namesSugar:
		scaling=True
	if ("3" in resname and atname=="O3\'") or ("5" in resname and atname=="O5\'"): #dont scale terminal hydroxyl groups
		scaling=False
	return scaling


# check if all arguments were given, if topology file exists, if scaling factor is a valid number, if mask is not provided in 1-5 format and if parmed is available. Exit if problem
intro="\nthis is script to STAFIX-scale amber topology with parmed\nusage:\n\tpython scaleSTAFIX.py parm_file scaling_factor [mask]\n\n\t\t[mask] is to be given in format resid1,resid2,resid3,... (e.g. 1,2,3,4). If no mask is specified, all RNA residues will be scaled\n"
try:
	inputTOP=sys.argv[1]
	if not os.path.exists(inputTOP):
		print("\n!invalid input file!")
		raise
	scalefact=float(sys.argv[2])
	try:
		mask=sys.argv[3]
	except:
		mask="*"
	if "-" in mask:
		raise
	if not find_executable("parmed"):
		print("\n!cannot execute parmed!")
		raise 
except:
	print(intro)
	sys.exit()

inputTOPfilemane,inputTOPextension=os.path.splitext(inputTOP)
new_parm=inputTOPfilemane+"STAFIX"+str(scalefact)+inputTOPextension
TMPfile="TMPstafixParmed1.out"
parmed_scale_input="TMPstafixParmed2.in"
present_atom_types={}
#definition of RNA residue names and of atom names for atoms that are NOT to be scaled
residue_namesA="A","A3","A5","RA","RA3","RA5"
residue_namesG="G","G3","G5","RG","RG3","RG5"
residue_namesC="C","C3","C5","RC","RC3","RC5"
residue_namesU="U","U3","U5","RU","RU3","RU5"
atom_namesSugar="O2\'","HO2\'","HO3\'","HO5\'"
atom_namesA="N3","N6","H61","H62"
atom_namesG="N1","H1","N2","H21","H22","N3"
atom_namesC="O2","N4","H41","H42"
atom_namesU="O2","N3","H3"

# remove pamed out file. if it exists, parmed does not override it 
if os.path.exists(TMPfile):
	os.system("rm "+TMPfile)

#run parmed for the first time to get info about present atoms:
os.system("parmed "+inputTOP+"<< EOF >> "+TMPfile+"\nprintDetails :"+mask+"\nEOF") 

#load TMPstafixParmed1.out file and read info about atom types present and store it into dictionary present_atom_types. Dictionary format is: {atomtype : [atomids, LJradius, LJepsilon]}
with open(TMPfile, "r") as TMPfile1:
	for line in TMPfile1:
		if "ATOM" in line:
			break
	for line in TMPfile1:
		if not line.strip():      
			break
		else:
			data=line.split()
			atomid,resname,atname,atomtype,ljrad,ljdepth=data[0],data[2],data[3],data[4],float(data[6]),float(data[7])
			if is_to_be_scaled(resname,atname):
				if atomtype not in present_atom_types:	#add new atom type to dictionary
					present_atom_types[atomtype]=["@"+atomid,ljrad,ljdepth]
				else:					#add the atomid to the existing atom type
					masklist=present_atom_types[atomtype][0]
					present_atom_types[atomtype]=[masklist+","+atomid,ljrad,ljdepth]

#remove new_parm file. if it exists, parmed does not override it
if os.path.exists(new_parm):
	os.system("rm "+str(new_parm))

# make parmed input with definition of new atom types and changeLJpair for those atoms, that are to be scaled.
with open(parmed_scale_input,"w") as parmedIn:
	for atom_type_group in present_atom_types:
		ljrad,ljdepth=present_atom_types[atom_type_group][1],present_atom_types[atom_type_group][2]
		parmedIn.write("addLJType "+present_atom_types[atom_type_group][0]+" radius "+str(ljrad)+" epsilon "+str(ljdepth)+"\n")

	pat_tuple=list(present_atom_types.items())
	for i in range (0,len(pat_tuple)):
		for j in range (0,i+1):
			group1,group2=pat_tuple[i][1][0],pat_tuple[j][1][0]
			eps1,eps2=pat_tuple[i][1][2],pat_tuple[j][1][2]
			eps_orig=sqrt(eps1*eps2)
			eps=str(eps_orig*scalefact)
			r=str(pat_tuple[i][1][1]+pat_tuple[j][1][1])
			parmedIn.write("changeLJPair "+group1+" "+group2+" "+r+" "+eps+"\n")

	parmedIn.write("parmout "+new_parm+"\ngo")

# run parmed and clean
os.system("parmed "+inputTOP+"< "+parmed_scale_input)
os.system("rm "+TMPfile+" "+parmed_scale_input)

#print "how to cite STAFIX usage" ??

if mask=="*":
	mask="all RNA residues"
print(("\nnew file generated: " +new_parm+"\nscaled residues: "+mask+"\n"))
	
